# Exploiting an Epigenetic Resistance Mechanism to PI3 Kinase Inhibition in Leukemic Stem Cells

**DOI:** 10.1101/2025.07.11.663968

**Authors:** Shira G. Glushakow-Smith, Imit Kaur, Simone Sidoli, Shayda Hemmati, Ellen Angeles, Taneisha Sinclair, Samarpana Chakraborty, Aaliyah Battle, Kristina Ames, Swathi-Rao Narayanagari, Rotila Hyka, Mark Soto, Melissa Tracy, Jayaram Vankudoth, Seiya Kitamura, Linde Miles, Ulrich Steidl, Aditi Shastri, Amit Verma, Kira Gritsman

## Abstract

Acquired non-genetic resistance mechanisms to existing therapies contribute to poor outcomes for acute myeloid leukemia (AML) patients, and inability to target leukemic stem cells (LSCs) can lead to relapse. To overcome these challenges, we tested whether LSCs have dependencies on PI3 kinase (PI3K). We found that LSCs are susceptible to isoform-selective targeting of PI3K and are particularly dependent on the P110 alpha isoform of PI3K. We discovered that PI3K inactivation leads to dynamic changes in EZH2/PRC2 function in leukemic cells, and we uncovered downregulation of EZH2 protein levels as a resistance mechanism in response to PI3K inhibition. We found that PI3K inhibition in AML cells can lead to compensatory upregulation of EZH1, and that EZH1 knockdown can sensitize AML cells to PI3K inhibition. We leveraged this resistance mechanism by combining a PI3K inhibitor with an EZH1/2 dual inhibitor, which successfully overcomes the acquired resistance and leads to sustained targeting of AML cells ex vivo and in murine AML and PDX models in vivo. This study identifies a promising novel therapeutic regimen for targeting LSCs in AML.

## Introduction

Acute Myeloid Leukemia (AML) is a devastating disease for which the prognosis has not improved significantly over the last 10 years. This is due to the high toxicity of currently available treatments and the fact that 50-70% of AML patients will relapse after achieving complete remission (1). One reason for this is that most current therapeutic approaches for AML are focused on targeting the highly proliferative leukemic blasts, which make up the bulk of the disease. However, this often leaves behind the leukemic stem cells (LSCs) that can contribute to minimal residual disease (MRD), which is associated with relapse and poor prognosis in AML (2). Both LSCs and blasts often develop resistance to standard AML treatments. Resistance to cancer therapeutics may arise from genetic mechanisms, such as selective pressure for specific mutations. However, a less well understood mechanism of resistance to therapy involves non-genetic mechanisms, which include epigenetic changes or transcriptional plasticity as a result of therapeutic pressure(3).

AML is molecularly heterogeneous, with a variety of cytogenetic alterations and driver mutations(4). While targeted therapeutics for some mutations are available, this heterogeneity makes it difficult to develop more universal therapeutics for AML, and emergence of resistant clones with different mutations is common. One of the two most common classes of driver mutations occurs in activated signaling molecules, such as FLT3, C-KIT, or RAS. Activation of the PI3K/AKT pathway, which is downstream of all of these signaling molecules, is observed in up to 80% of AML cases(5), making this pathway an attractive target for more universal therapeutic approaches. PI3K and AKT are rarely themselves mutated in AML, but AKT is frequently phosphorylated, including in LSCs (6–8). Additionally, we and others have shown that activation of AKT in mouse hematopoietic cells can drive both AML and T-ALL (9–11).

Class I PI3Ks are a family of heterodimeric lipid kinases composed of a regulatory subunit and a catalytic subunit (12). These kinases are responsible for directly activating signal transduction pathways via recruitment to the plasma membrane by receptor tyrosine kinases, as well as other phosphorylated proteins. This begins a signaling cascade in which phosphatidyl inositol phosphate-2 (PIP2) is converted to PIP3, and the subsequent accumulation of PIP3 allows for the signal transmission into the cell and activation of downstream effectors such as AKT, a serine/threonine kinase that regulates many key processes in the cell (12, 13). PI3K/AKT pathway inhibition has been demonstrated to inhibit the proliferation of AML cells in preclinical studies (14–17). However, despite multiple early phase clinical trials (18), no PI3K or AKT inhibitors have been approved for AML treatment to date. This could be due to the inability to achieve sufficient potency in AML cells without excessive toxicity, as the PI3K/AKT pathway is also required for many normal processes in healthy cells. Furthermore, therapeutic resistance could have been an issue, and potential resistance mechanisms to PI3K inhibition in AML are poorly understood.

In the hematopoietic compartment, PI3K signaling is important for maintaining HSC homeostasis (9–11, 19), so it is important to determine a safe way to target this pathway in leukemia patients. There are three catalytic subunits of Class 1A PI3K (P110α, β, and δ), encoded by *Pik3ca*, *Pik3cb*, and *Pik3cd*, respectively. These isoforms can all bind to the same regulatory subunits p85α and p85β, allowing for some redundancy between them. To determine if isoform-selective inhibition could be a safe and effective strategy in AML, we previously generated conditional knockout models for individual Class IA PI3K isoforms (P110α or P110β) in mouse hematopoietic cells and in leukemic cells (20, 21). We found that myeloid leukemias can have isoform-selective dependencies on PI3K (20, 21). Recent findings have also shown leukemic dependency on Class 1B PI3K (P110γ) in AML (22–24). In other forms of cancer, isoform-selective inhibition of PI3K has been shown to be an effective therapeutic strategy, and several isoform-selective inhibitors have been approved by the FDA for use in patients with chronic lymphocytic leukemia, lymphoma, or breast cancer (25).

The second most common class of driver mutations in AML is in epigenetic regulators, such as DNA methyltransferases and histone methyltransferases (4), which are observed in over 75% of AMLs. Among epigenetic regulators implicated in AML, the histone methyltransferase Enhancer of Zeste Homolog 2 (EZH2) has been widely studied not only in AML, but also in other cancers. EZH2 is a core catalytic component of the Polycomb Repressive Complex 2 (PRC2), which deposits the repressive H3K27me3 mark. EZH2 can be directly mutated or impacted by mutations in other proteins that either disrupt or enhance its function(26, 27). It can function as either a tumor suppressor or oncogene, depending on the context(27). EZH1, a closely related PRC2-associated methyltransferase with partially redundant function to EZH2, has been shown to compensate for EZH2 loss in leukemic cells (28, 29).

To determine a safe way to target the PI3K pathway in LSCs, we asked whether LSCs in AML could have isoform-selective PI3K dependencies. Here we report that LSCs are selectively dependent on the P110α isoform of PI3K, and that PI3K inactivation, either genetically or pharmacologically, promotes myeloid differentiation and loss of self-renewal. Our studies also uncovered a non-genetic mechanism of resistance to PI3K inactivation - downregulation of EZH2 protein levels, which makes resistant cells more dependent upon EZH1. We found that the combination of a PI3K inhibitor with an EZH1/2 inhibitor can effectively leverage this acquired resistance mechanism to deplete the LSC pool in AML.

## Methods

### Sex As a Biological Variable

Our study examined both male and female animals and samples from both male and female patients, and similar findings are reported for both sexes.

### Mice

Mice were maintained under pathogen-free conditions in a barrier facility in microisolator cages based under protocols #00001165 and #00001181, which were approved by the Institutional Animal Care and Use Committee at Albert Einstein College of Medicine (AECOM). *Pik3cd* germline KO, *Pik3ca*-lox/lox, and *Pik3cb*-lox/lox were described previously(30). For *Pik3ca*-lox/lox;*Pik3cb*-lox/lox excision, pIpC (Sigma-Aldrich) was dissolved in Hanks’ balanced salt solution, and 250 μg was injected intraperitoneally three times on nonconsecutive days. For transplantation experiments, donor mice were 6-10 weeks old and recipient mice were 6-8 weeks old. Genotypes of each allele (*Pik3ca*, *Pik3cb*, and *Pik3cd*) were determined by PCR using genomic DNA from tails as previously described(30).

### Generation of Leukemic Mice

To generate MLL-AF9 leukemic mice, Lineage-Sca1+cKit+ cells were sorted into RPMI media with 10ng/ml IL3, 10ng/ml IL-6, 50ng/ml SCF, 10ng/ml TPO and 20ng/ml Flt3L (Peprotech). Cells were transduced with viral sup generated from the plasmid MSCV-KMT2A-MLLT3-GFP. Transduced cells were plated in methylcellulose for 7 days. 150,000-200,000 sorted GFP+ cells and 250,000 CD45.1+ B6.SJL (The Jackson Laboratory; Strain No. 002014) helper bone marrow cells were transplanted into lethally irradiated (9 Gy) C57Bl/6 (Taconic) recipient mice. To generate the NPM1c/NRAS leukemia model, 250,000-300,000 Npm1^LSL-CA/+;^Nras^LSL-G12D^ frozen bone marrow cells (provided as a gift from Dr. Linde Miles) and 700,000 CD45.1 + B6.SJL helper bone marrow cells were transplanted into lethally-irradiated C57/Bl6 recipient mice. Patient-derived xenotransplantation was performed by transplanting 1,200,000 cells from frozen bone marrow from Patient sample #7 (Supplementary Table 1) into sub-lethally irradiated (2 Gy) NSG mice. Donor cells were injected into the tail-vein and recipient mice were given 100 mg/mL Baytril-100 (Bayer) in drinking water for 4 weeks after transplantation. Mice were euthanized upon signs of development of AML.

### Bone Marrow Aspiration

For femoral bone marrow aspiration mice were anesthetized. The animal’s leg was first disinfected with 70% ethanol, a fine needle was then inserted into the femoral bone marrow cavity through the distal condyles. A small volume (up to 10 uL) of bone marrow was aspirated as this has been shown not to compromise the animal’s functionality of the leg or their overall health. Sides (left/right) were alternately used for sequential aspirates. The animals received 5mg/kg Banamine preemptive per subcutaneous injection as an analgesic.

### Drug Treatment of Mice

The mice were treated 6mg/kg copanlisib three times per week on alternating days intraperitoneally, and with 100mg/kg valemetostat five days a week by oral gavage, or with a combination of both drugs. The control mice were treated with solvents of respective drugs: copanlisib vehicle: PEG400/acidified water, pH 3.5-4 intraperitoneally (3 times per week) or valemetostat vehicle: 0.5% MC Solution by oral gavage 5 times per week.

### Cell culture

NOMO1, MOLM13, MOLM14, and HEL cells were cultured in RPMI with 10% fetal bovine serum (FBS) and 1% Penicillin-Streptomycin (P/S). The venetoclax resistant cell line MOLM13-VR was generated as previously described (31). OCI-AML3 cells were cultured in α-MEM with 20% FBS and 1%P/S. All cells were kept in a 5% CO_2_ 37°C incubator. Cell lines were authenticated with STR profiling with ATCC and DSMZ databases by the Albert Einstein genomics core facility and verified to be free of mycoplasma on a monthly basis.

### Plasmids

The MSCV-KMT2A-MLLT3-GFP and MSCV-KMT2A-MLLT3-NEO plasmids were a gift from Scott Armstrong’s lab. pcDNA3-3myc-6His-EZH2 21A and pcDNA3-3myc-6His-EZH2 21D were a gift from Mien-Chie Hung (Addgene plasmids # 42663 and # 42664). The cDNA from each EZH2 mutant plasmid was subcloned into pMSCV-IRES-mCherry FP, which was a gift from Dario Vignali (Addgene plasmid # 52114). MSCV-EZH2-PGK-Puro-IRES-GFP was a gift from Christopher Vakoc (Addgene plasmid # 75125). The Cre-ER puro plasmid was a gift from the Gilliland lab.

### Colony Forming Assays

Cells were plated in 35×10mm dishes of Mouse Methylcellulose Complete Media (R&D HSC007). For ex vivo excision of PI3K alleles in MSCV-Cre-ER-PURO transduced colonies, a 4-Hydroxytamoxifen (4-OHT) stock was made in ethanol, and then dissolved in HSC007 methylcellulose at a final concentration of 20nM. Dishes were incubated at 37C for seven days. Colonies were manually counted under an inverted microscope (Olympus CKX41). Cells were then washed off the plate with PBS. A portion of the cells were prepared for microscopy using Cytofuge2 Cytocentrifuge (StatSpin) cytospin and stained with Wright-Giemsa stain (Fisher Scientific 22-122911). Higher resolution images were acquired using a Zeiss Axiovert 200M Microscope with a digital camera. Image acquisition was performed using AxioVision software.

### Human Patient Sample Assays

Healthy human CD34+ BM cells were purchased from Stem Cell Technologies and stored at - 150C. Experiments with patient samples were approved under the Albert Einstein College of Medicine Institutional Review Board IRB protocol #2015-4600. De-identified patient samples were obtained with written informed consent through the Albert Einstein College of Medicine biobank IRB protocol #2005-536, in accordance with the Declaration of Helsinki. All patient and BM donor characteristics are listed in Table S1. The samples were defrosted in RPMI 1640, washed, resuspended in RPMI 1640/FBS (fetal bovine serum) media (20% FBS, 1% penicillin-streptomycin) and cultured for 1 hour at 37C. Cells were passed through a 30-μm cell strainer to remove dead cell clumps, washed with RPMI 1640, and resuspended in IMDM +2%FBS media. Cells were plated in quadruplicate in MethoCult H4435 Enriched (StemCell Technologies) at single cell density at indicated concentrations ranging from 1-5×10^4^ cells/plate. Colonies were counted manually after 10-12 days in culture.

### Microarray Analysis and GSEA

The GFP+ GMPs were sorted from the bone marrow of transplant recipient mice following the gating strategy in Fig S3D using the Aria II instrument (BD Biosciences). Cells were sorted into 50μl RNA extraction buffer (ARCTURUS PicoPure RNA Isolation Kit (Life Technologies, Invitrogen) and stored at −80°C. RNA was isolated using the (ARCTURUS PicoPure RNA Isolation Kit (Life Technologies, Invitrogen) according to the manufacturer’s instructions. RNA quantification was performed using the RNA Quantification Kit for SYBR Green I and ROX™ Passive Reference Dye (Thermo Fisher, catalog no. 902905). Total RNA was amplified and hybridized to the Gene Chip ® Mouse Transcriptome Pico Assay 1.0 (Affymetrix part number: 902663) using the GeneChip™ Hybridization, Wash, and Stain Kit (catalog no. 900720). Raw data was analyzed for quality control using Expression Console software (Affymetrix). After quality control, microarray data was analyzed by Gene Set Enrichment Analysis (GSEA) using MSigDB software (http://software.broadinstitute.org/gsea/index.jsp), Transcriptome Analysis Software (Affymetrix), and Gene Ontology Enrichment analysis (Gene Ontology Consortium; geneontology.org). The microarray data is available at the GEO Expression Omnibus under accession number GSE261355.

### Cell Proliferation Assays

Cells were plated in four 96-well plates (1 x 10^5^ cells/well) and treated with drugs in triplicate. Drugs and media were replenished every 3 days. At 0 hours, 24 hours, 48 hours, 72 hours, 6 days, 9 days, and 12 days the amount of ATP present was measured using CellTiter-Glo Luminescent Cell Viability Assay (Promega, Catalog no. G7570). Cell Titer Glo reagent was added 1:1 before allowing the plate to gently rock 20 minutes at room temperature. Luminescence measured by a PerkinElmer Victor X5 Multilabel Plate Reader.

### Flow Cytometry

For flow cytometry analysis of stem and progenitor cell populations, bone marrow cells were stained with a lineage cocktail of species-appropriate biotin-labeled lineage antibodies for 30 min at 4C, followed by species appropriate fluorochrome-conjugated surface antibodies for 20 min at 4C (Table S2-4). For non-stem and progenitor flow analysis, cells were stained with species-appropriate fluorochrome-conjugated surface antibodies for 20 min at 4C (Table S2-4). For intracellular flow analysis, cells were stained with all surface antibodies as described above, and then fixed with cytofix/cytoperm solution (BD biosciences BDB554714) according to the manufacturers protocol. Cells were then stained with either fluorochrome-conjugated intracellular antibodies or unconjugated primary antibodies (Table S3) for 30 min at room temperature, followed by species appropriate fluorochrome-conjugated secondary antibody for 30 min at room temperature. Flow sorting was performed on BD FACSAria III. Flow cytometry analysis was performed on the BD FACS LSRII or Cytek Aurora. Cell cycle analysis was performed with Ki-67 and Hoechst staining. Apoptosis analysis was performed using an APC Annexin V with 7-AAD kit (BD Biosciences), following the manufacturers protocol. Analysis of all flow cytometry data was performed using FlowJo software (version 10).

### Time Course Drug Treatment

MOLM14 or NOMO1 cells were treated with 1μM alpelisib (BYL719; S2814 Selleckchem), 1 μM buparlisib (BKM120; S2247 Selleckchem), 100nM or 500nM Copanlisib (BAY 90-6946; S2802 Selleckchem) or DMSO control and harvested at 3 hour, 24 hour, and 48 hour time points. Cell pellets were stored at −80°C.

### Western Blotting

Protein was extracted from frozen cells with Pierce IP Lysis Buffer and Halt Protease/Phosphatase Inhibitor Cocktail (Thermo Fisher), 30uL/10^6^ cells, after PBS wash. Sonication was used to extract nuclear proteins. Lysate Protein concentration was determined using a Bradford assay with Bio-Rad Protein Assay Dye Reagent. Proteins were separated with SDS-PAGE on a NuPAGE Bis-Tris 10% polyacrylamide gel (Thermo Fisher) or a Novex Tris-Glycine 10-20% polyacrylamide gel and transferred to a 0.2um nitrocellulose membrane. The membranes were blocked in Tris-Buffered Saline 0.1% Tween 20 (TBST) with 5% bovine serum albumin (BSA) and washed with TBST. The membranes were incubated overnight at 4°C with primary antibodies as listed in Table S5. Internal loading controls were incubated 1 hour at room temperature. After washing, the membrane was blotted with fluorescent secondary antibodies for 1 hour. Bands were visualized using the Odyssey Fc imaging system (LI-COR) and protein band quantification was measured using Image Studio (version 5.2.5). Please refer to Table S5 for all Western antibody information. Some membranes were stripped with ReBlot Plus Strong antibody stripping solution (Millipore #2504) and then re-blocked and re-probed.

### RT-PCR

RNA was isolated from frozen cell pellets using the Qiagen RNeasy and Qiashredder kits per manufacturer’s instructions. cDNA was made from isolated RNA using RNA to cDNA EcoDry Premix (Random Hexamers) (Takara #639546) as per manufacturer’s instructions. PCR was performed using SYBR-Green reagents and acquired on the Viia7 Real-Time PCR system. For primer sequences see Table S6.

### Histone extraction and digestion

Histone proteins were extracted from the pellet as described by (32) to ensure good-quality identification and quantification of single histone marks. Briefly, histones were acid-extracted with chilled 0.2 M sulfuric acid (5:1, sulfuric acid : pellet) and incubated with constant rotation for 4 h at 4°C, followed by precipitation with 33% trichloroacetic acid (TCA) overnight at 4°C. Then, the supernatant was removed and the tubes were rinsed with ice-cold acetone containing 0.1% HCl, centrifuged and rinsed again using 100% ice-cold acetone. After the final centrifugation, the supernatant was discarded and the pellet was dried using a vacuum centrifuge. The pellet was dissolved in 50 mM ammonium bicarbonate, pH 8.0, and histones were subjected to derivatization using 5 µL of propionic anhydride and 14 µL of ammonium hydroxide (all Sigma Aldrich) to balance the pH at 8.0. The mixture was incubated for 15 min and the procedure was repeated. Histones were then digested with 1 µg of sequencing grade trypsin (Promega) diluted in 50mM ammonium bicarbonate (1:20, enzyme:sample) overnight at room temperature. Derivatization reaction was repeated to derivatize peptide N-termini. The samples were dried in a vacuum centrifuge.

### Sample desalting

Prior to mass spectrometry analysis, samples were desalted using a 96-well plate filter (Orochem) packed with 1 mg of Oasis HLB C-18 resin (Waters). Briefly, the samples were resuspended in 100 µl of 0.1% TFA and loaded onto the HLB resin, which was previously equilibrated using 100 µl of the same buffer. After washing with 100 µl of 0.1% TFA, the samples were eluted with a buffer containing 70 µl of 60% acetonitrile and 0.1% TFA and then dried in a vacuum centrifuge.

### LC-MS/MS Acquisition and Analysis

Samples were resuspended in 10 µl of 0.1% TFA and loaded onto a Dionex RSLC Ultimate 300 (Thermo Scientific), coupled online with an Orbitrap Fusion Lumos (Thermo Scientific). Chromatographic separation was performed with a two-column system, consisting of a C-18 trap cartridge (300 µm ID, 5 mm length) and a picofrit analytical column (75 µm ID, 25 cm length) packed in-house with reversed-phase Repro-Sil Pur C18-AQ 3 µm resin. Peptides were separated using a 30 min gradient from 1-30% buffer B (buffer A: 0.1% formic acid, buffer B: 80% acetonitrile + 0.1% formic acid) at a flow rate of 300 nl/min. The mass spectrometer was set to acquire spectra in a data-independent acquisition (DIA) mode. Briefly, the full MS scan was set to 300-1100 m/z in the orbitrap with a resolution of 120,000 (at 200 m/z) and an AGC target of 5×10e5. MS/MS was performed in the orbitrap with sequential isolation windows of 50 m/z with an AGC target of 2×10e5 and an HCD collision energy of 30.

Histone peptides raw files were imported into EpiProfile 2.0 software (33). From the extracted ion chromatogram, the area under the curve was obtained and used to estimate the abundance of each peptide. In order to achieve the relative abundance of post-translational modifications (PTMs), the sum of all different modified forms of a histone peptide was considered as 100% and the area of the particular peptide was divided by the total area for that histone peptide in all of its modified forms. The relative ratio of two isobaric forms was estimated by averaging the ratio for each fragment ion with different mass between the two species. The resulting peptide lists generated by EpiProfile were exported to Microsoft Excel and further processed for a detailed analysis.

Proteomics data submitted to ProteomeXchange via the PRIDE database, accession #: PXD050834.

### Valemetostat Synthesis

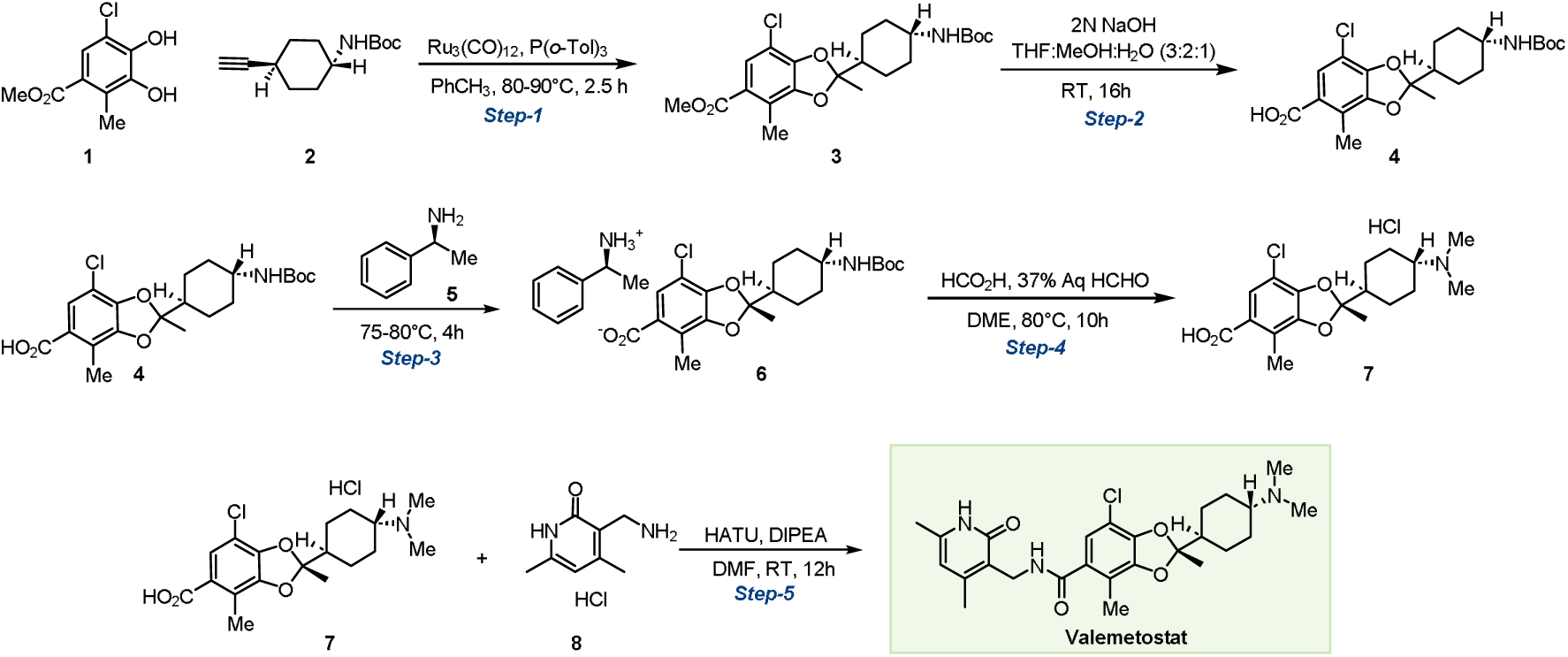

The synthetic procedure was adapted from patent WO/2022/009911.

**Methyl2-((1*r*,4*r*)-4-((*tert*-butoxycarbonyl)amino)cyclohexyl)-7-chloro-2,4-dimethylbenzo[*d*][1,3]dioxole-5-carboxylate** (**3**). In a pear-shaped flask equipped with a condenser and a PTFE-coated stirring bar, a mixture of methyl 5-chloro-3,4-dihydroxy-2-methylbenzoate **1** (8 g, 1.0 equiv., 0.04 mol, Accela, cat. # SY291378), tri-*o*-tolylphosphine (3 g, 0.3 equiv., 0.33 mol, 1PlusChem, cat. # 1P003689) and triruthenium dodecacarbonyl (2 g, 0.1 equiv., 4 mmol 1PlusChem, cat. #1P0039U2) was purged with argon/vacuum cycles (4 cycles), then toluene (84 mL) was added. The system was purged with four additional argon/vacuum cycles, and the reaction was then stirred at 120° C for 30 minutes. A mixture of *tert*-butyl (1*r*,4*r*)-4-ethynylcyclohexyl) carbamate **2** (9 g, 1.2 equiv., 0.04 mol, 1PlusChem, cat. #1P00IIXA) in toluene (70 mL) was added to the dark mixture, and the resulting orange solution was stirred for another 2 hours at 120° C. Upon reaction completion (confirmed by TLC), the solvent was evaporated under rotary evaporation, and the crude residue was purified by flash column chromatography (Teledyne CombiFlash Nextgen 300+, SiliaSep PREMIUM cartridge, 10 to 15 % EtOAc in hexane) to afford compound **3** as a pale green semi-solid (13.5 g, 68%). **^1^H NMR** (300 MHz, CDCl_3_) δ 7.53 (s, 1H), 4.37 (s, 1H), 3.84 (s, 3H), 3.39 (s, 1H), 2.38 (s, 3H), 2.07 (dd, *J* = 9.3, 5.6 Hz, 2H), 1.99 – 1.92 (m, 2H), 1.90 – 1.76 (m, 1H), 1.62 (s, 3H), 1.43 (s, 9H), 1.38 – 1.28 (m, 2H), 1.18 – 1.02 (m, 2H).

**2-((1*r*,4*r*)-4-((*tert*-Butoxycarbonyl)amino)cyclohexyl)-7-chloro-2,4-dimethylbenzo[*d*][1,3]dioxole-5-carboxylic acid** (**4**). In a round-bottom flask equipped with a PTFE-coated stirring bar, compound **3** (13 g,1.0 equiv., 30 mmol) was dissolved in THF (106 mL) and MeOH (56 mL), then 2M aqueous NaOH (35 mL) was added. The reaction was stirred at room temperature for 16 hours. Upon completion (confirmed by TLC), the organic solvent was removed by rotary evaporation, and the crude was treated with 2N HCl to adjust pH to 2-3. Water (200 mL) was added to the reaction mixture, causing precipitation of copious solid, which was collected by filtration over a Buckner funnel, washed with 250 mL of hexane), and dried overnight under vacuum compound **4** as an off-white solid (11 g, 87%). **^1^H NMR** (300 MHz, CDCl_3_) δ 7.59 (s, 1H), 4.42 (s, 1H), 3.39 (s, 1H), 2.38 (s, 3H), 2.07 (d, *J* = 11.9 Hz, 2H), 2.00 – 1.91 (m, 2H), 1.83 (tt, *J* = 12.1, 3.2 Hz, 1H), 1.61 (s, 3H), 1.44 (s, 9H), 1.30 (t, *J* = 12.5 Hz, 2H), 1.10 (q, *J* = 12.3 Hz, 2H). **MS**: *m/z* calculated for: C_21_H_28_ClNO_6_ [M+H-Boc]^+^: 325.79 found: 325.79,

**(*S*)-1-Phenylethan-1-aminium(*R*)-2-((1*r*,4*R*)-4-((*tert*-butoxycarbonyl)amino)cyclohexyl)-7-chloro-2,4-dimethylbenzo[*d*][1,3]dioxole-5-carboxylate** (**6**).

*<u>Step 1.</u>* In a round-bottom flask equipped with a PTFE-coated stirring bar, **4** (10.5 g, 1 equiv., 24.7 mmol) in 105 mL of 1,2-dimethoxyethane (DME) was stirred at 75-80 °C. (*S*)-1-Phenylethan-1-amine **5** (3.6 mL, 1.13 equiv., 27.9 mmol, Sigma-Aldrich, cat. # 8070470010) was added in one portion, then the reaction mixture was stirred at 80 °C for 4 hours. A mixture of DME (47 mL) and H_2_O (17 mL) was heated to 60 °C was added to the reaction mixture, and the reaction was allowed to cool to room temperature, causing precipitation of a solid. The precipitate was filtered out and washed with DME (200 mL), then dried under vacuum to obtain partially enriched compound **6** as an off-white solid (3.5 g).

*<u>Step 2.</u>* In a round-bottom flask equipped with a PTFE-coated stirring bar, **6** (3.5 g) was added to a mixture of DME (47 mL) and water (16 mL). To the solution, 1.12 mL of 5M HCl was added at room temperature. After 10 minutes, the reaction mixture was heated to 75 °C, and a solution of (*S*)-1-phenylethan-1-amine (0.72 mL, Sigma-Aldrich, cat. # 8070470010) in DME (5.5 mL) was added dropwise over 10 minutes. The reaction was stirred at 80 °C for 2 hours, then cooled 0 °C, causing the precipitation of a solid. The solid was collected by filtration and washed with DME (100 mL) to afford enriched compound **6** as an off-white solid (2.6 g, 25% over 2 steps). **^1^H NMR** (600 MHz, CD_3_OD) δ 7.47 – 7.38 (m, 5H), 7.09 (s, 1H), 4.41 (q, *J* = 6.9 Hz, 1H), 3.29 – 3.24 (m, 1H), 2.28 (s, 3H), 1.96 (tq, *J* = 9.0, 3.3 Hz, 4H), 1.83 (tt, *J* = 12.0, 3.0 Hz, 1H), 1.61 (d, *J* = 6.8 Hz, 3H), 1.59 (s, 3H), 1.43 (s, 9H), 1.35 – 1.27 (m, 2H), 1.23 – 1.15 (m, 2H). The ^1^H NMR spectrum closely matches the literature report (WO/2022/009911)

**(*R*)-7-Chloro-2-((1*r*,4*R*)-4-(dimethylamino)cyclohexyl)-2,4-dimethylbenzo[*d*][1,3]dioxole-5-carboxylic acid hydrochloride** (**7**). In a round-bottom flask equipped with a PTFE-coated stirring bar, to a stirred solution of **6** (2.53 g, 1.0 equiv., 5.90 mmol) in DME (2.53 mL) was added formic acid (5.04 mL, 22.5 equiv., 134.00 mmol, Fluka, cat. #60-006-16) and 37% aqueous formaldehyde (3.75 mL, 17.1 equiv., 102.00 mmol, Thermo-Fisher, cat. #119690010) under a nitrogen atmosphere. The reaction mixture was stirred at 75-80 °C for 10 hours, then upon completion (confirmed by LC-MS) the solvent was removed under rotary evaporation to afford the crude product, which was purified by reverse-phase flash column chromatography (Teledyne CombiFlash Nextgen 300+, SiliaSep PREMIUM C18 cartridge - 40 g, 100% MeCN + 0.01% formic acid to 100% H_2_O + 0.01% formic acid), the compound eluted in 20% MeCN in H_2_O. Fractions containing the product were dried by rotary evaporation and then lyophilized to afford the **7** as an off-white solid (1.4 g, 67%) **^1^H NMR** (600 MHz, CD_3_OD) δ 7.37 (s, 1H), 3.20 (tt, *J* = 12.2, 2.6 Hz, 1H), 2.83 (s, 6H), 2.35 (s, 3H), 2.19 – 2.10 (m, 4H), 2.04 – 1.97 (m, 1H), 1.66 (s, 3H), 1.56 (tt, *J* = 12.9, 6.6 Hz, 2H), 1.42 (tdd, *J* = 16.0, 7.3, 4.0 Hz, 2H). **MS**: *m/z* calculated for: C_18_H_25_ClNO_4_ [M+H]^+^: 354.15 found: 354.10.

**Valemetostat.** In a round-bottom flask equipped with a PTFE-coated stirring bar, **7** (1.25 g, 1.0 equiv., 3.53 mmol), **8** (1.33 g, 2.0 equiv., 7.10 mmol, 1pluschem, cat #: 1P000EJW) and HATU (2.68 g, 2.0 equiv., 7.01 mmol, 1PlusChem, cat. # 1P001LPW) were dissolved in DMF (12 mL), then DIPEA (6.31 mL, 10.0 equiv., 35.00 mmol, Sigma-Aldrich, cat. # D125806) was added under argon atmosphere, and the reaction mixture was stirred at room temperature for 10 hours. Upon completion of the reaction (confirmed by LC-MS), the reaction mixture was basified with 0.1 N aqueous NaOH to pH 10-11, then the reaction mixture was extracted three times with EtOAc (200 mL each time), the combined organic layers were dried over Na_2_SO_4_ and concentrated under rotary evaporation. The obtained crude was dissolved in 150 mL of water and stirred overnight, causing the precipitation of a solid which was collected by filtration and dried under vacuum.The obtained solid was washed once with 1M aqueous NaHCO_3_ (200 mL) and solid-extracted three times with DCM: MeOH (9:1, 200 mL each time). The organic layer was dried over Na_2_SO_4_ and concentrated under rotary evaporation, then the obtained compound was dissolved in a minimal amount of MeCN:H_2_O (3:7) and lyophilized to afford the valemetostat as a white solid (1.28 g, 74.2%) **^1^H NMR** (600 MHz, DMSO-*d_6_*) δ 11.47 (s, 1H), 8.12 (t, *J* = 5.0 Hz, 1H), 6.85 (s, 1H), 5.86 (s, 1H), 4.22 (d, *J* = 5.0 Hz, 2H), 3.32 (s, 2H), 2.24 – 2.01 (m, 16H), 1.92 – 1.78 (m, 5H), 1.60 (s, 3H), 1.22 – 1.10 (m, 4H). **^13^C NMR** (151 MHz, DMSO*-d_6_*) δ 167.03, 163.43, 149.93, 147.35, 144.01, 143.23, 132.03, 123.23, 121.98, 121.35, 116.25, 109.02, 107.81, 63.04, 45.87, 41.71, 35.51, 27.52, 25.59, 25.57, 22.40, 19.36, 18.65, 12.45. **MS**: *m/z* calculated for: C_26_H_34_ClN_3_O_4_ [M+H]+:488.03. found: 488.30, Purity (HPLC-UV 254 nm): >95%. (*t*_R_= 3.91 min).

The following method was used to check the enantiomer purity:

Column: Daicel, CHIRALCEL OZ-H, 4.6 mm ID X 250 mm L

Elution solvent: n-hexane: ethanol: diethylamine = 60:40:0.04 (v/v)

Flow rate = 1 mL/min, HPLC, detection at 254 nm

Temperature: 25°C, the major isomer eluted at t_R=_ 5.97 min and its retention time closely matches the literature report (US20170073335A1).

### Reagents

Please refer to Tables S2-5 for the list of Western blot antibodies and flow cytometry antibodies used. All other reagents are listed in their respective methods section.

### Study approval

All mouse breeding and animal experiments were approved by the Institutional Animal Care and Use Committee under protocol nos. 20170205, 20170206, 00001165, and 00001181.

Experiments with patient samples were approved under IRB#2015-4600 and #2005-536.

### Statistics

GraphPad Prism 10 was used for all statistical analysis. For the comparison of two experimental groups, an unpaired t-test was used. For the comparison of more than two groups, ANOVA test was used with Tukey’s multiple comparisons test. In all graphs, error bars indicate mean ± SEM. A *P* value less than 0.05 was considered significant (*), with 0.01 (**), 0.001 (***), and 0.0001 (****) representing higher levels of significance.

## Results

### PI3K inactivation impairs leukemic stem cell self-renewal and promotes myeloid differentiation

To study the effects of targeting PI3K in LSCs, we performed colony forming assays on sorted Lin-Sca1+c-Kit+ (LSK) cells from *Pik3ca*^lox/lox^, *Pik3cb*^lox/lox^, *Pik3cd*^-/-^, *Pik3ca* ^lox/lox^;*Pik3cb*^lox/lox^ , or *Pik3ca* ^lox/lox^;*Pik3cd*^-/-^ mice transduced with the KMT2A-MLLT3-NEO (MLL-AF9-NEO) retrovirus (Figure 1, Supplementary Figure S1), which can generate an aggressive AML when introduced into hematopoietic stem and progenitor cells and confers serial replating ability(34). After 4 weeks of serial replating, we acutely excised the PI3K alleles by introducing a Cre-ER-puro retrovirus, selecting transduced cells with puromycin, and then inducing excision of floxed alleles with 4-orthohydroxytamoxifen (4-OHT) (Figure 1A). We found that deletion of *Pik3ca*, *Pik3cb*, or pairs of isoforms from leukemic stem cells caused a significant reduction in the serial replating capacity of KMT2A-MLLT3 cells (Figure 1B). However, while transduction of *Pik3cd*^-/-^ LSK cells with KMT2A-MLLT3 initially generated fewer colonies than WT cells, *Pik3cd* deletion alone did not abrogate serial replating (Supplementary Figure S1). Together, these data suggest that leukemic stem cell self-renewal is selectively dependent on specific Class I PI3K isoforms.

**Figure 1:**
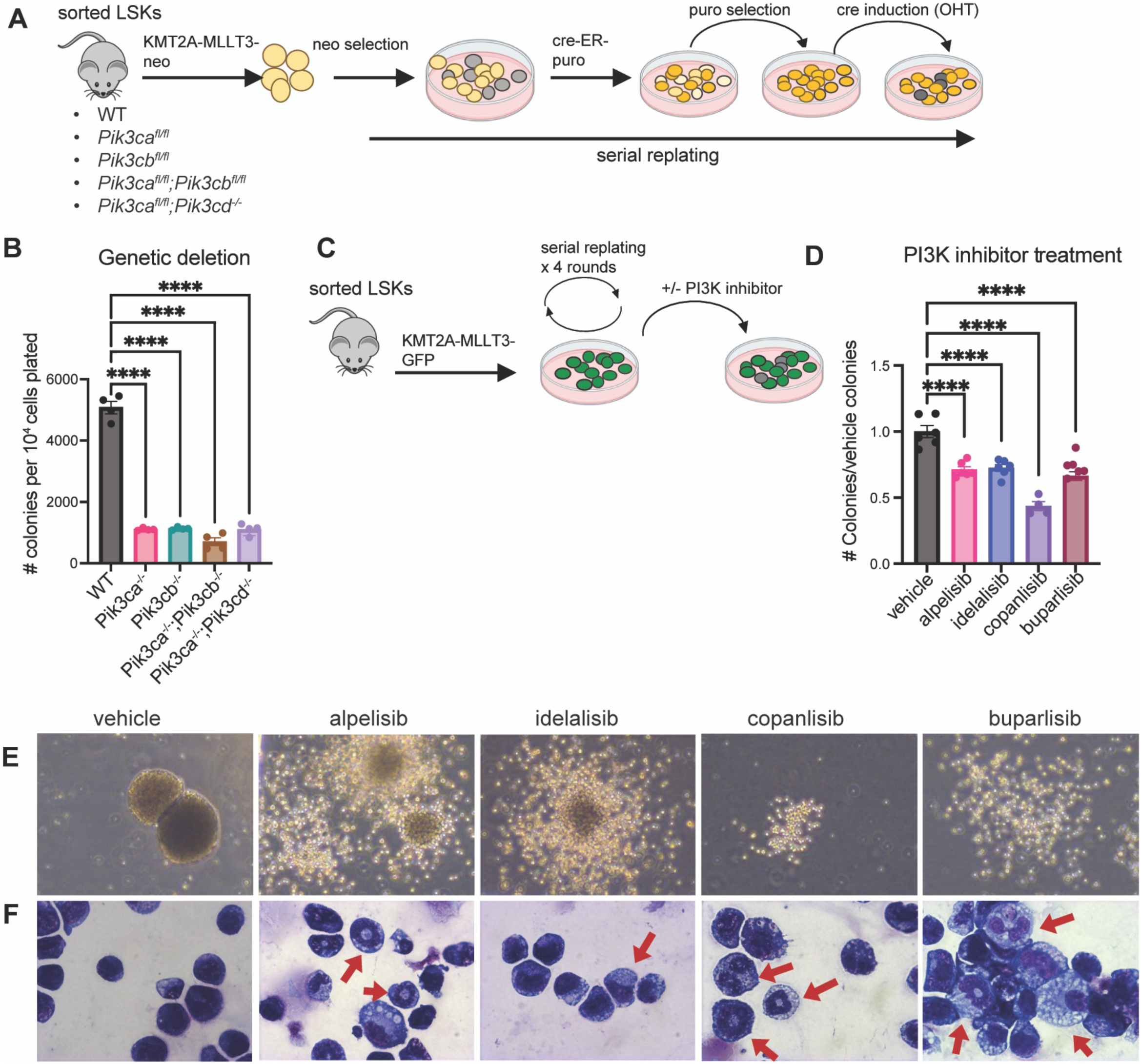
PI3K Disruption Impairs Leukemic Stem Cell Self-Renewal and Promotes Myeloid Differentiation. **(A)** Experimental schematic for generation of PI3K KO LSC colonies with inducible excision of PI3K isoforms **(B)** KMT2A-MLLT3 leukemic stem cell colonies after 4 weeks of serial re-plating at one week after acute excision of PI3K isoforms (n=4) **(C-F)** Serial replating of KMT2A-MLLT3 leukemic stem cell colonies in the presence of PI3K inhibitors (alpelisib = a selective, idelalisib = d selective, copanlisib = a/d selective, TGX221 = b selective, buparlisib = pan-PI3K inhibitor) **(C)** Experimental schematic **(D)** Colony counts normalized to vehicle control **(E)** Representative colony images captured using 400x objective **(F)** Representative images of cytospins made from colonies captured using 630x objective. Red arrows indicate cells with monocytic or neutrophilic morphologies. Each experiment was performed at least 3 times. Each value is presented as mean +/- standard error of the mean (SEM). One-way ANOVA test with Tukey’s multiple comparisons was used in B and D. ****P < 0.0001

We next examined the effects of pharmacologic inhibition of PI3K on LSCs by performing serial replating assays on WT LSK cells transduced with the KMT2A-MLLT3-Neo retrovirus in the presence of either isoform-selective PI3K inhibitors (P110α: alpelisib(35), P110β: TGX221(36), P110δ:idelalisib(37), P110α/δ: copanlisib(38)), or the pan-PI3K inhibitor buparlisib(39) (Figure 1C). We observed that pharmacologic inhibition of individual PI3K isoforms was sufficient to significantly impair self-renewal of KMT2A-MLLT3 cells (Figure 1D). Interestingly, the morphology of KMT2A-MLLT3 colonies indicated signs of myeloid differentiation upon treatment with PI3K inhibitors, which was corroborated by morphologic analysis of individual cells from the colonies, showing some features of monocytes or neutrophils (Figure 1E, F). Consistent with these findings in mouse LSCs, the PI3K inhibitor copanlisib also induced myeloid differentiation in the human AML cell lines MOLM14, NOMO1 and OCI-AML3, as evidenced by changes in morphology and increased expression of the monocytic markers CD15 and CD11b (Supplementary Figure S2). This suggests that PI3K inhibition not only impairs LSC self-renewal but can also promote myeloid differentiation.

### PI3K deletion depletes functional leukemic stem cells *in vivo*

We next wanted to analyze the impact of targeting PI3Kα or PI3Kβ, which were required for serial replating, in leukemic progression *in vivo*. We injected sorted LSK cells from *Pik3ca*^lox/lox^;Mx1-Cre or *Pik3cb*^lox/lox^;Mx1-Cre mice transduced with the KMT2A-MLLT3-GFP retrovirus into lethally irradiated C57Bl/6 recipients (Figure 2A). We found that deletion of *Pik3cb* in leukemic mice had no significant effect on survival or disease progression (Supplementary Figure S3A). Additionally, deletion of *Pik3cb* had no effect on leukemia initiating cell (LIC) frequency as tested by secondary transplantation (Supplementary Figure S3B). We also did not observe any significant changes in spleen weight, though liver weights were slightly decreased in *Pik3cb*^-/-^ mice (Supplementary Figure S3C). We confirmed by PCR that the leukemic cells in *Pik3cb*^-/-^ mice did have excision of exon 2 of *Pik3cb* as expected (Supplementary Figure S3D)(40). This suggests that PI3Kβ does not play an important role in KMT2A-MLLT3 AML *in vivo*.

**Figure 2:**
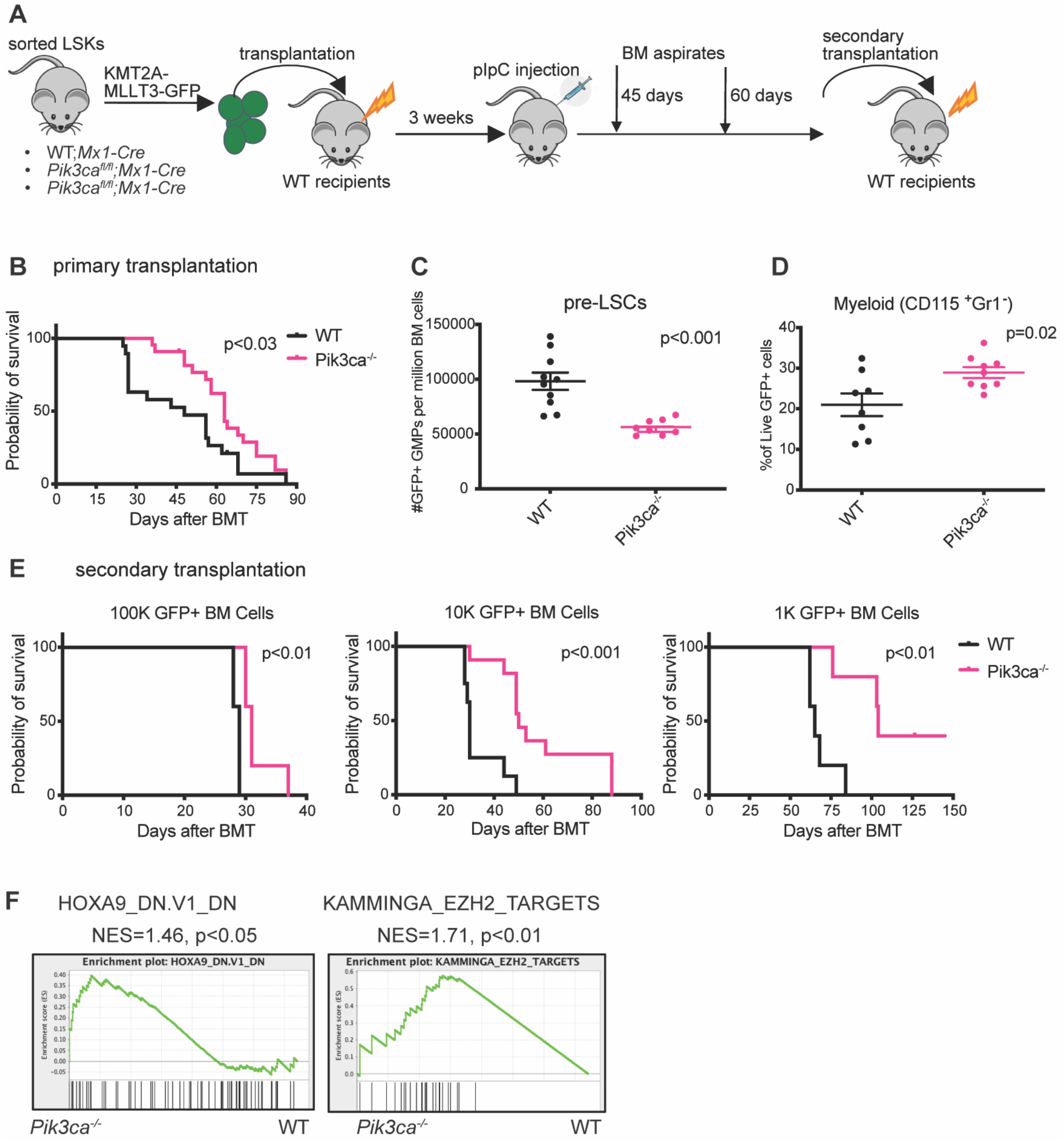
PI3K Deletion Impairs Leukemic Stem Cell Function in vivo. **(A)** Experimental schematic for KMT2A-MLLT3 retroviral transduction and transplantation assay with acute excision of PI3K isoforms in recipient mice **(B)** Kaplan-Meier survival curve from primary transplantation. Log-rank analysis was used. (n = 19-22 mice per group) **(C)** Flow cytometry analysis of GFP+ GMPs from 45-day bone marrow (BM) aspirates **(D)** Flow cytometry analysis of 60-day BM aspirates gated on GFP+ cells. (n=8-10 per group). Each value is presented as mean +/- standard error of the mean (SEM). Unpaired t-test was used in C and D. **(E)** Kaplan-Meier survival curves from secondary BM transplantation (BMT) with limiting numbers of GFP+ leukemic cells transplanted (n= 5-10 mice per group). Log-rank analysis was used. Each experiment was performed at least three times with similar results **(F)** Gene set enrichment (GSEA) plots of microarray data from pre-leukemic LSCs (GFP+ GMPs) sorted from pre-clinical BM aspirates

In contrast, deletion of *Pik3ca* significantly prolonged survival in the KMT2A-MLLT3 group compared to WT controls (Figure 2B). To follow disease evolution, we performed bone marrow aspiration on transplanted mice at two pre-clinical stages, before they showed clinical signs of disease. At the earliest pre-clinical time point, *Pik3ca* deletion resulted in a decrease in the number or immunophenotypically defined LSCs (Figure 2C), defined as GFP+ granulocyte-macrophage progenitors for the KMT2A-MLLT3 AML model (Supplementary Figure S4A) (41). At the second pre-clinical time point, we also observed an increase in expression of the monocytic marker CD115 in bone marrow aspirates from the *Pik3ca*^-/-^ group (Figure 2D). This data suggests that *Pik3ca* plays an important role in maintaining the self-renewal capacity of LSCs *in vivo*. To determine whether *Pik3ca* is also required to maintain functional leukemia-initiating cells (LICs), we performed secondary transplantation of decreasing doses of GFP+ leukemic cells from the primary transplant recipients into sub-lethally irradiated mice. We found that at each dose of leukemic cells transplanted, survival was significantly prolonged in the recipients of *Pik3ca*-deleted leukemic cells, consistent with a depletion in functional LICs (Figure 2E). This suggests that PI3Kα is required to sustain functional LIC activity *in vivo*.

### Dynamic changes in EZH2 regulation contribute to AML resistance to PI3K inhibition

To better understand the mechanism through which *Pik3ca* deletion leads to a reduction in functional LSCs, we sorted GFP+ GMPs from preclinical bone marrow aspirates of KMT2A-MLLT3-GFP primary transplant mice (Supplementary Figure S4) and performed DNA microarray analysis in triplicate using the Affymetrix Mouse Transcriptome Assay 1.0 kit. Gene Set Enrichment Analysis (42, 43) of *Pik3ca*^-/-^ pre-LSCs compared with WT pre-LSCs revealed downregulation of several AKT and AKT-MTOR gene signatures, as expected (Supplementary Figure S5A). This GSEA also revealed significant enrichment of a HOXA9 knockdown signature (HOXA9_DN.V1_DN) in *Pik3ca*^-/-^ pre-LSCs, consistent with loss of self-renewal (Supplementary Figure S5A, Figure 2F). Interestingly, we also observed enrichment of the KAMMINGA_EZH2_TARGETS signature (Supplementary Figure 5B, Figure 2F), suggesting altered regulation of PRC2-associated transcriptional programs in *Pik3ca*^-/-^ pre-LSCs.

Although *Pik3ca* deletion prolonged survival in the KMT2A-MLLT3 AML model, mice in the *Pik3ca* – deleted group eventually still developed leukemia with a similar phenotype as control WT;Mx1-Cre mice (Figure 2B, Supplementary Figure S6A-C), suggesting the emergence of resistance mechanisms. To test for potential adaptive resistance mechanisms to targeting PI3K in leukemic cells, we performed Western analysis on bone marrow cells from moribund leukemic primary transplant recipient mice. First, to determine whether other isoforms of PI3K are compensating for loss of *Pik3ca,* we examined AKT phosphorylation at Ser473. We observed a complete loss of AKT phosphorylation in *Pik3ca^-/-^* leukemic cells (Figure 3A), indicating that compensation by other PI3K isoforms is not a likely mechanism of resistance. Given our transcriptome data showing potential effects of *Pik3ca* deletion on EZH2, we also assessed EZH2 protein levels. We found that total EZH2 protein levels were also decreased in *Pik3ca*^-/-^ leukemic cells compared to WT, indicating that a loss of EZH2 protein may be a potential resistance mechanism to targeting PI3K (Figure 3A). However, we could still detect H3K27me3 in *Pik3ca*-/- leukemic cells (Figure 3A).

**Figure 3:**
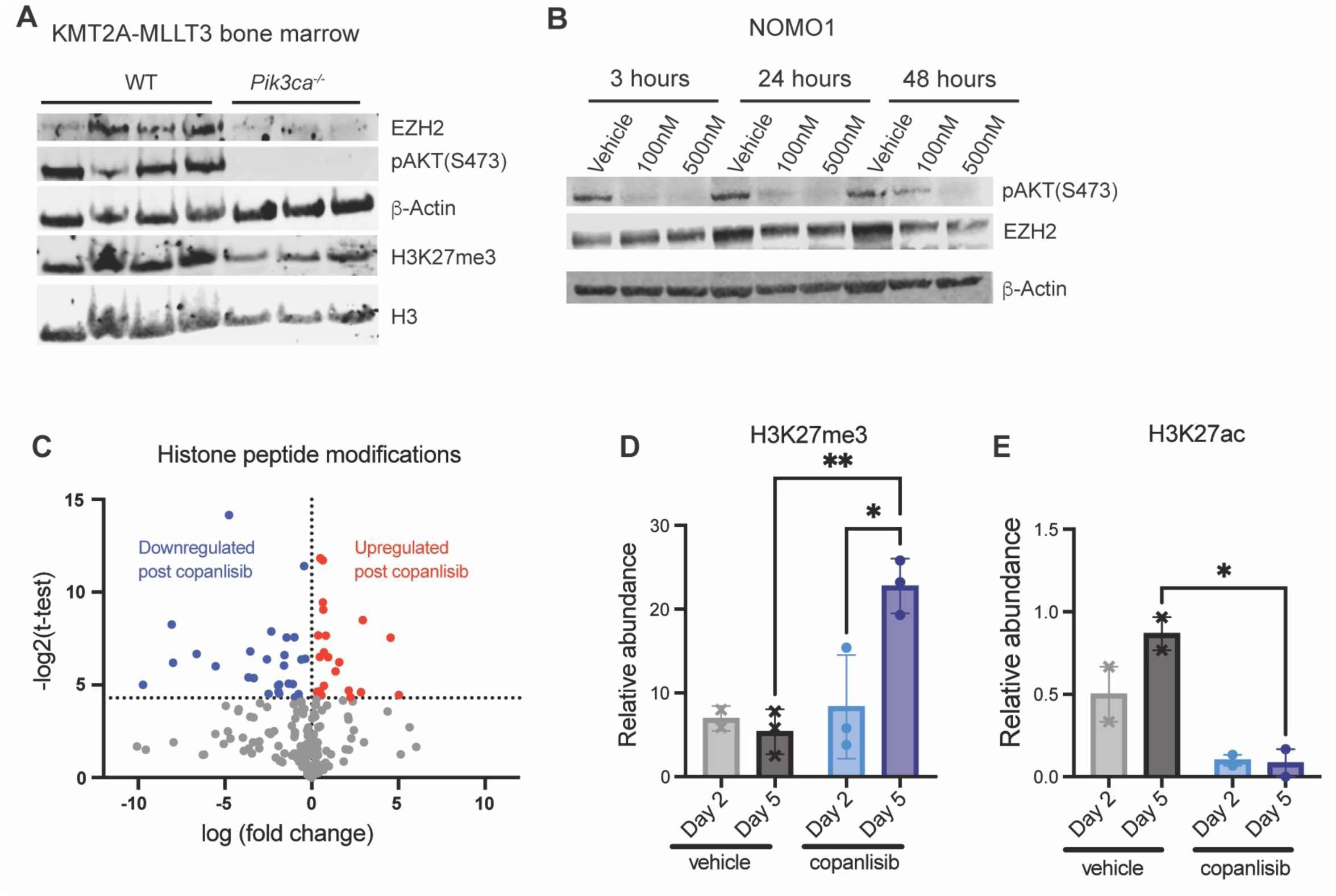
Dynamic Changes in EZH2 Contribute to AML Resistance to PI3K Inhibition. **(A)** Western Blot analysis on bone marrow cells from moribund primary KMT2A-MLLT3 transplant recipient leukemic mice **(B)** Western blot analysis on NOMO1 cells treated for 2 days with 2 doses of copanlisib. **(C)** Volcano plot highlighting the changes in histone peptide modifications in NOMO1 cells after 5 days of treatment with 500nM copanlisib **(D)** Relative abundance of the H3K27me3 mark in NOMO1 cells treated with 500nM copanlisib **(E)** Relative abundance of the H3K27Ac mark in NOMO1 cells treated with 500nM copanlisib. Each value is presented as mean +/- standard error of the mean (SEM). (n = 3 per group) One-way ANOVA test with Tukey’s multiple comparisons was used in E and F. **P < 0.01 *P < 0.05. These experiments were performed two times with similar results.

To determine if we could recapitulate this decrease in EZH2 protein in response to PI3K inactivation in AML cell lines, we treated NOMO1 and MOLM14 cells with copanlisib over the course of two days. Western analysis revealed a decrease in total EZH2 protein levels only after 24 hours of exposure to the PI3K inhibitor, with the greatest decrease observed within 48 hours (Figure 3B), suggesting an adaptive response rather than an acute effect. To functionally examine the results of EZH2/PRC2 regulation as a result of PI3K inhibition, we performed histone proteomics on NOMO1 cells after treatment with copanlisib for 2 or 5 days. We found an increasing number of significantly dysregulated histone peptide modifications after prolonged exposure to PI3K inhibitors (Figure 3C). Surprisingly, we observed a significant increase in H3K27me3 relative abundance after 5 days of copanlisib treatment and a corresponding decrease in abundance of the H3K27ac activating mark, despite the decrease in EZH2 protein levels (Figure 3D, E). This suggests compensatory regulation of PRC2-mediated chromatin states during the adaptive resistance response to PI3K inhibition.

### PI3K inhibition cooperates with EZH1/2 dual inhibition to target leukemic cells

Previous studies have shown that when EZH2 is compromised, its homologue EZH1, another histone methyltransferase, becomes essential for leukemic cell function (28, 29). To determine if EZH1 can compensate for loss of EZH2 after PI3K inhibition to maintain H3K27me3, we examined the expression of EZH1 in copanlisib-treated AML cell lines by qPCR. We observed a time-dependent upregulation of EZH1 RNA expression in both NOMO1 and MOLM14 cells after copanlisib treatment (Figure 4A). We also confirmed sustained EZH1 protein expression in MOLM14 cells after 48 hours of copanlisib treatment (Supplementary Figure S7A). To determine if EZH1 can compensate for EZH2 loss in the setting of PI3K inhibition in AML cells, we performed shRNA knockdown of EZH1 in NOMO1 cells. Using two different shRNAs targeting EZH1, we generated stable NOMO1 cells with EZH1 knockdown, as confirmed with qPCR (Figure 4B). Proliferation assays revealed that EZH1 knockdown can sensitize NOMO1 cells to copanlisib (Figure 4B). Importantly, we found that knockdown of EZH1 had no impact on cell proliferation at baseline (Supplementary Figure S7B). However, the shEZH1 cells were more sensitive to copanlisib in a dose dependent manner (Figure 4C, Supplementary Figure S7B), indicating a context-specific dependency on EZH1 following PI3K inhibition.

**Figure 4:**
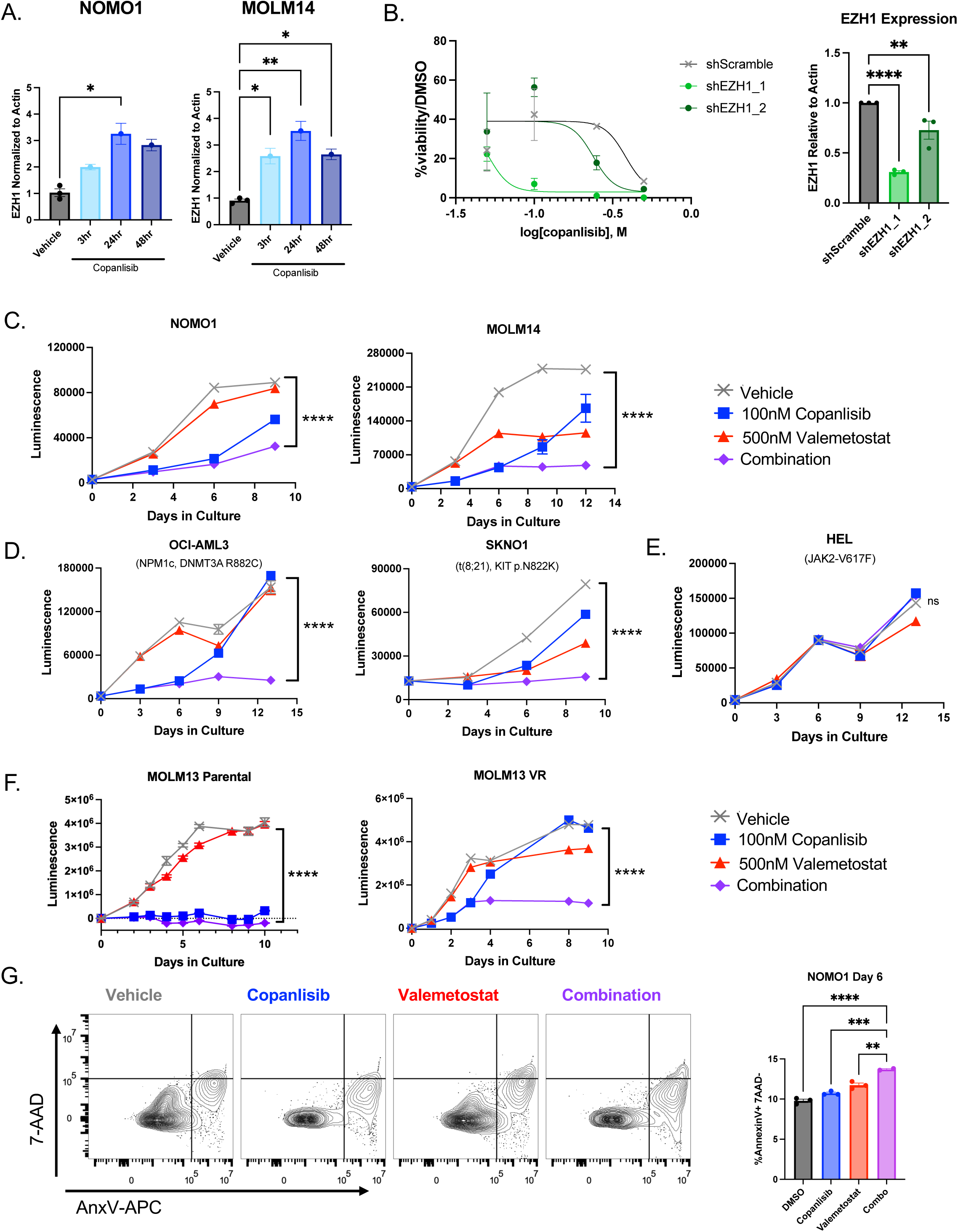
PI3K Inhibitors Cooperate with EZH1/2 Dual Inhibition to Target LSCs. **(A)** qPCR testing EZH1 expression levels in NOMO1 and MOLM14 cells treated with 200nM Copanlisib over the course of 2 days (n = 3) **(B)** Dose response curve on day 11 of treatment of NOMO1 shEZH1 lines with copanlisib treatment. qPCR showing knockdown of EZH1 expression in NOMO1 shEZH1 cells is on the right (n= 3) **(C-F)** Proliferation assays with copanlisib +/- valemetostat treatment of KMT2A-rearranged AML cell lines **(C)** and non-KMT2A-rearranged AML cell lines **(D,E)**, and MOLM13 parental and venetoclax resistant AML cell lines **(F).** Each experiment was performed at least three times with similar results. **(G)** Representative gating strategy for apoptotic cells in NOMO1 cells after 6 days of treatment (left). Quantification of apoptotic populations (right, n = 3). Each experiment was performed at least two times with similar results. Each value is presented at mean +/- standard error of the mean (SEM). One-way ANOVA test with Tukey’s multiple comparisons was used in A, B, and F. ****P ≤ 0.0001 ***P ≤ 0.001 **P<0.01 *P ≤ 0.05 ns = not significant.

We hypothesized that pharmacologic inhibition of EZH1 could overcome the resistance to PI3K inhibition that we observed in AML cells driven by the loss of EZH2 and lead to synthetic lethality in PI3K inhibitor-treated cells. Therefore, we tested the EZH1/2 dual inhibitor valemetostat, which has been shown to have preclinical activity in AML(44, 45), in combination with copanlisib in Cell Titer Glo proliferation assays in multiple AML cell lines. Our data suggest that in most cases, copanlisib alone could initially suppress proliferation, but that within a week of treatment proliferation increased again (Figure 4C,D). However, when copanlisib was combined with valemetostat, proliferation was suppressed for the duration of the experiment (Figure 4C,D). Importantly, this was observed in a variety of AML cell lines with diverse molecular and cytogenetic profiles, both KMT2A-rearranged (Figure 4C) and not KMT2A-rearranged (Figure 4D). However, we observed resistance to this combination treatment in HEL cells, indicating that the anti-proliferative effects of this drug combination are not due to nonspecific cytotoxic effects (Figure 4E).

Because venetoclax resistance is becoming a major clinical problem in AML treatment, we also tested our drug combination in the venetoclax-resistant cell line MOLM-13VR (46). We found that while parental MOLM13 cells are sensitive to copanlisib alone and to the combination, MOLM-13VR cells become resistant to copanlisib but are still sensitive to the copanlisib and valemetostat combination treatment (Figure 4F).

To determine how this combination treatment impacts proliferation of leukemic cells, we examined myeloid differentiation, cell cycling, and apoptosis by flow cytometry. First, we tested whether the combination treatment promotes more terminal differentiation of leukemic cells than PI3K inhibition alone. Flow cytometry analysis showed no significant differences in myeloid marker expression with combination drug treatment compared to the PI3K inhibitor alone, so the combination effects of our treatment are unlikely to be a result of differentiation effects (Supplementary Figure S8A, B). Next, we performed cell cycle analysis using Hoechst 33342 vs Ki-67. After 48 hours of treatment, we observed an increase in the proportion of leukemic cells in G0 cell cycle arrest in our combination treated group compared to either single agent or the vehicle control in both NOMO1 and OCI-AML3 cells (Supplementary Figure S8C,D). Cytospins of leukemic cells treated for six days showed morphological changes consistent with cell death (Supplementary Figure S8E).

This led us to perform apoptosis assays with Annexin V and 7-AAD to determine if our combination treatment induces synthetic lethality. We found that after six days of treatment, our combination-treated cells exhibited significantly higher levels of apoptosis in NOMO1 cells compared to either single agent or the vehicle control (Figure 4G). This suggests that the combination of copanlisib and valemetostat induces cell cycle arrest and apoptosis, consistent with synthetic lethality of this drug combination in AML cells.

### The PI3K inhibitor copanlisib cooperates with the EZH1/2 dual inhibitor valemetostat to target LSCs *in vivo*

To confirm our results with the copanlisib and valemetostat combination *in vivo*, we injected bone marrow cells from CD45.2 NPM1^LSL-CA/+;^NRAS^LSL-G12D^ leukemic mice(47) into lethally irradiated CD45.1 WT recipients. Once engraftment was confirmed, we randomized the mice by %CD45.2 in the peripheral blood and then treated them for four weeks with 6mg/kg copanlisib intraperitoneally, 100mg/kg valemetostat by oral gavage, or the combination (Figure 5A). The valemetostat used *in vivo* was synthesized by the Albert Einstein College of Medicine Chemical Synthesis Core Facility and was confirmed to reduce H3K27me3 in MOLM14 cells by Western blot (Supplementary Figure S9A) and to reduce proliferation in MOLM14 cells, alone and in combination with copanlisib (Supplementary Figure S9B). After four weeks of treatment with copanlisib and valemetostat, we continued to monitor any remaining animals for survival and then concluded the experiment at 8 weeks post treatment (Figure 5A,B). Upon completion of the experiment at 8 weeks post treatment, a statistically significant survival benefit was observed in the combination treatment group, and half of the combination group mice survived the duration of the experiment, whereas there were no surviving mice in any of the other three treatment groups (Figure 5B).

**Figure 5:**
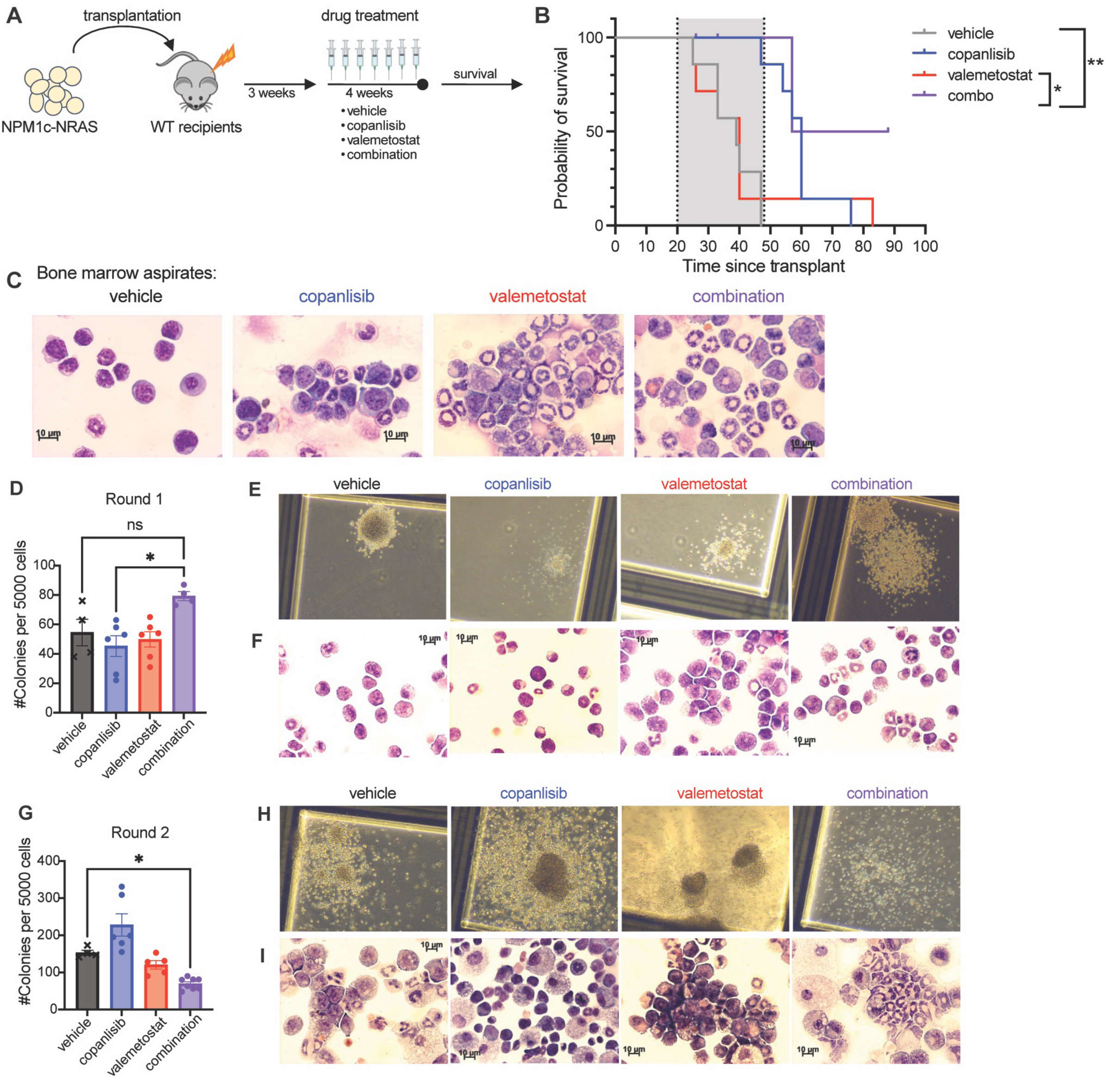
PI3K Inhibition Cooperates with EZH1/2 Dual Inhibition to Target LSCs in NPM1c-NRAS AML in vivo. **(A)** Experimental schematic for NPM1c-NRAS AML transplantation with *in vivo* drug treatment **(B)** Survival curves (n = 6-8 per group). The gray shaded area represents the duration of drug treatment. **(C)** Representative cytospin images made from Bone Marrow aspirates at 2 weeks post treatment start. **(D)** Results from Round 1 of re-plating assay using splenocytes from moribund mice.**(E-F)** Representative images of colonies **(E)** and colony cytospins **(F)** from Round 1 of re-plating. **(D)** Results from Round 2 of re-plating assay using splenocytes from moribund mice.**(G-I)** Representative images of colonies **(G)** and colony cytospins **(I)** from Round 2 of re-plating. Each value is presented at mean +/- standard error of the mean (SEM). One-way ANOVA test with Tukey’s multiple comparisons was used in D and F. *P < 0.05 ns = not significant. Each value is presented at mean +/- standard error of the mean (SEM). One-way ANOVA test with Tukey’s multiple comparisons was used in D and F. *P < 0.05 ns = not significant.

After two weeks of treatment, we performed pre-clinical bone marrow aspiration for analysis, to assess early cellular responses to drug treatment. Flow cytometry analysis showed no differences in disease burden, as demonstrated by similar levels of %CD45.2 across the treatment groups (Supplementary Figure S10A). Intracellular flow cytometry indicated that valemetostat is successfully able to achieve on target effects in immunophenotypically defined LSCs, resulting in a decrease in H3K27me3 levels (Supplementary Figure S9B,C). Cytospins of the bone marrow aspirates showed significant morphological changes in cells treated with valemetostat alone and in combination with copanlisib, consistent with myeloid differentiation (Figure 5C). To examine LSC function after drug treatment, we performed serial re-plating assays with splenocytes from drug-treated leukemic mice. We initially observed an overall increase in the number of colonies from combination treated mice (Figure 5D), and colony and cellular morphology were consistent with myeloid differentiation (Figure 5E,F). Upon re-plating these colonies, we found that cells from the combination-treated mice had decreased serial replating capacity, with a significant decrease in the number of colonies at Round 2 (Figure 5G). Furthermore, colonies from the combination group had morphological differences consistent with myeloid differentiation after serial replating (Figure 5H,I). Together, these data indicate a decrease in LSC self-renewal and an increase in myeloid differentiation *in vivo* upon combined copanlisib and valemetostat treatment.

Next, we sought to determine whether our combination regimen can impair leukemia initiating cell function *in vivo*. To test this, we injected sorted LSK cells transduced with the KMT2A-MLLT3-GFP retrovirus into lethally irradiated WT recipients. Once engraftment was confirmed, we randomized the mice by %GFP in the peripheral blood and then treated them for two weeks with 6mg/kg copanlisib intraperitoneally, 100mg/kg valemetostat by oral gavage, or the combination (Figure 6A). After two weeks of treatment, we euthanized the animals and performed disease burden analysis. In order to confirm on target effects of our drug treatments, we performed Western analysis on spleens from the primary transplant recipients. As expected, we observed a decrease in H3K27me3 in our valemetostat-treated and combination-treated mice, indicating that valemetostat was dosed appropriately (Supplementary Figure S11A,B). Given the aggressive nature of this model, we did not see any effect on spleen or liver weights or in the %GFP+ cells in the spleen or bone marrow after two weeks of treatment (Supplementary Figure S11D-E). However, we noted a significant increase in the frequency of the CD115+Gr1- cell population in each treatment group compared to the untreated controls, consistent with monocytic differentiation (Supplementary Figure S11F).

**Figure 6:**
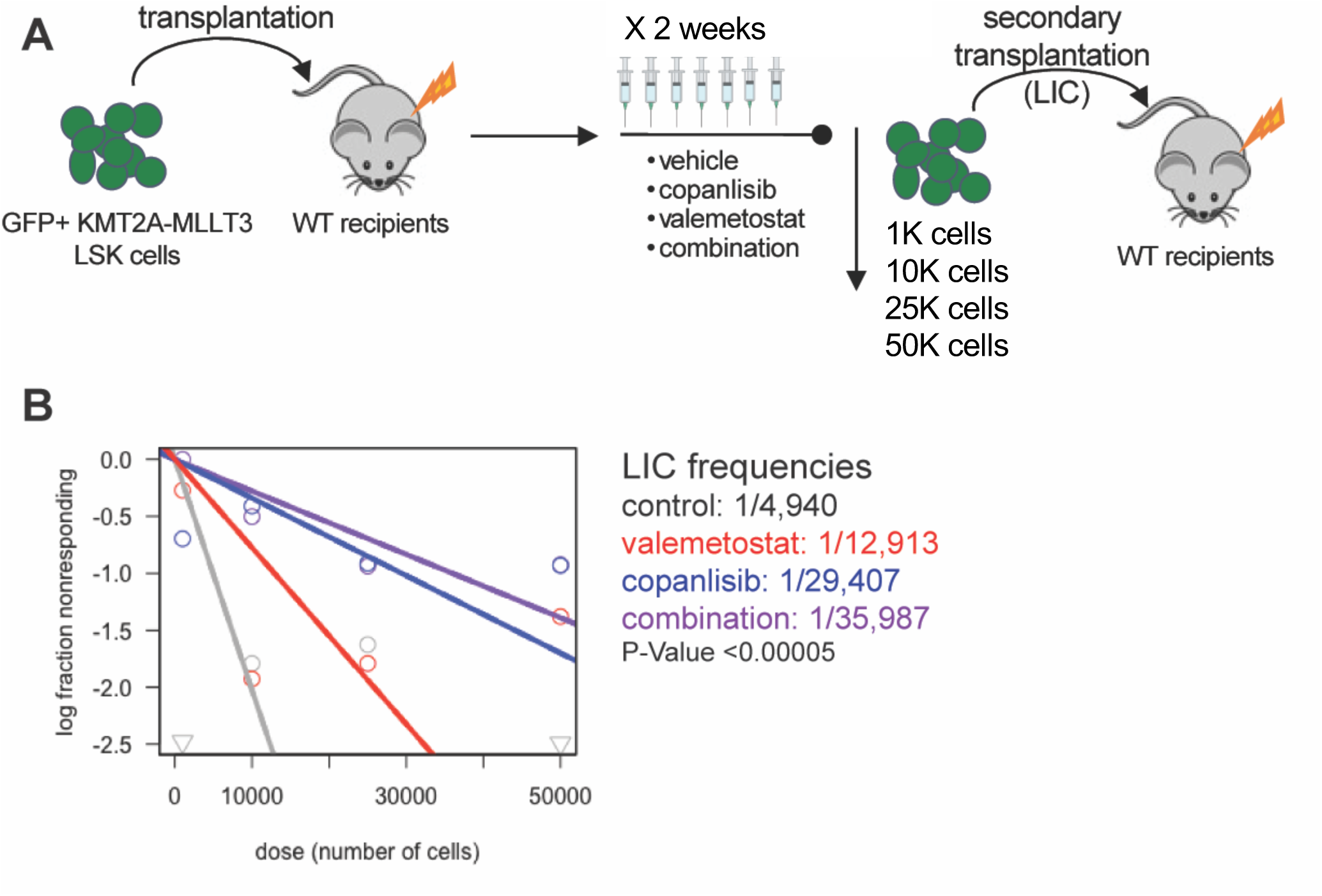
PI3K Inhibition Cooperates with EZH1/2 Dual Inhibition to Target LSCs in vivo. **(A)** Experimental schematic for bone marrow transplantation assay of KMT2A-MLLT3 AML cells with in vivo drug treatment with 6mg/kg copanlisib administered i.p. MWF +/- 100mg/kg valemetostat administered oral gavage, daily on M-F, followed by secondary transplantation LIC ELDA assay. **(B)** ELDA analysis based on survival data from secondary transplantation with limiting number of GFP+ leukemic cells.

To determine if this treatment can target leukemia initiating cells, we performed limiting dilution secondary transplantation with sorted GFP+ leukemic cells from mice in each treatment group, without any additional treatment in the secondary transplant recipients (Figure 6A). As expected, we observed prolonged survival in the combination group with transplantation of each cell dose compared to vehicle control, and at the lowest dose none of the combination treated mice developed leukemia during the observation period (Supplementary Figure S12). Importantly, extreme limiting dilution analysis (ELDA) (48) revealed a statistically significant 10-fold reduction in LIC frequency in our combination-treated mice compared to the untreated group (Figure 6B), demonstrating effective targeting of functional LICs with the copanlisib and valemetostat combination *in vivo*.

To determine the effects of the copanlisib and valemetostat combination on healthy hematopoietic cells, we treated whole bone marrow cells from WT mice in colony assays in methylcellulose supplemented with myeloid growth factors. Although we observed a statistically significant reduction in total colony numbers with our combination treatment (Supplementary Figure S13A), these cells were still able to produce every type of myeloid colony (Supplementary Figure S13B). This suggests the existence of a therapeutic window for combined PI3K and EZH1/2 targeting in the hematopoietic system.

### The copanlisib and valemetostat combination reduces colony formation by AML and MDS patient samples

To determine the clinical relevance of this drug combination in AML, we asked if this treatment regimen can also reduce the proliferation of AML and high-risk MDS patient cells. We tested our combination treatment in a primary MDS and AML patient samples with a variety of molecular subtypes (Supplementary Table 1) in methylcellulose colony assays with myeloid growth factors in the presence of copanlisib, valemetostat, or the combination. Consistent with our cell line data, we observed a significant reduction in the colony forming capacity in all of the patient samples we tested with the combination, compared to either single agent alone (Figure 7A-E; Supplementary Table 1).

**Figure 7:**
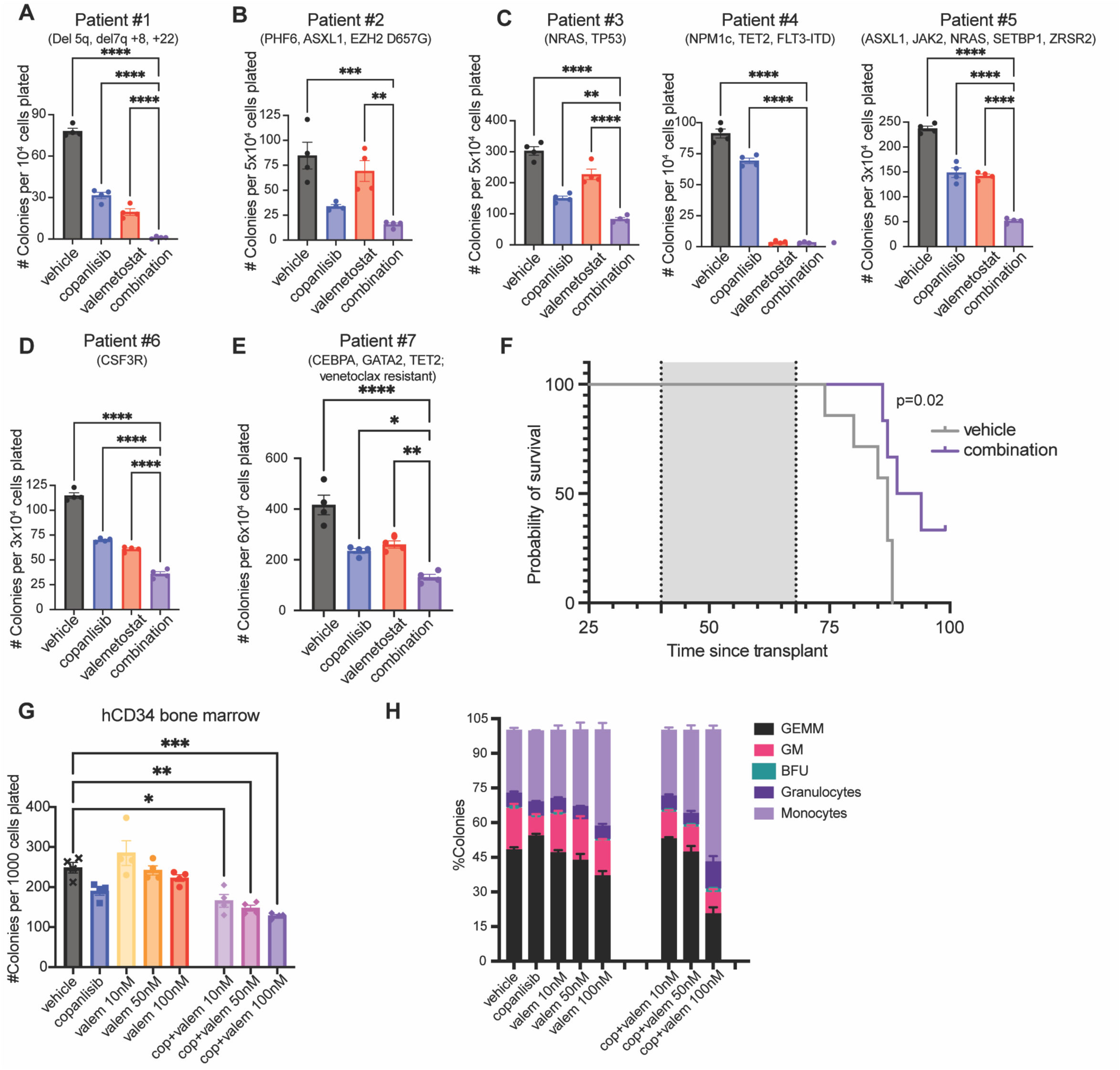
PI3K Inhibition Cooperates with EZH1/2 Dual Inhibition to Impair Colony Formation by AML and MDS Patient Cells. **(A)** Colony assays of AML patient sample #1 treated with 100nM copanlisib and 500nM valemetostat **(B)** Colony assay of AML patient sample #2 treated with 100nM copanlisib and 100nM valemetostat. **(C)** Colony assay of AML patient samples #3-5 treated with 100nM copanlisib and 50nM valemetostat. **(D)** Colony assay of AML patient sample #6 treated with 100nM copanlisib and 10nM valemetostat. **(E)** Colony assay of AML patient sample #7 treated with 100nM copanlisib and 50nM valemetostat **(F)** Kaplan-Meier survival curve of PDX using AML patient sample #7 comparing treatment groups. Log-Rank test was used. N=6 per group. **(G-H) (G)** Colony counts and **(H)** colony type breakdown on colony assays performed on hCD34+ cells treated with 100nM copanlisib in combination with valemetostat. Each value is presented as mean +/- standard error of the mean (SEM). One-way ANOVA test with Tukey’s multiple comparisons was used in A-E, and G. ****P < 0.0001 ***P < 0.001 **P < 0.01 *P < 0.05

Importantly, in several patient samples we observed effects of combination treatment on colony formation with lower doses of valemetostat (10-100nM) than the dose needed to observe a decrease in proliferation in AML cell lines (500nM) (Figure 7B-D), suggesting an increased sensitivity of primary patient cells to valemetostat. Furthermore, we observed activity of this drug combination even in difficult to treat AML subtypes, such as AML with complex cytogenetics (Figure 7A), with mutated TP53 (Figure 7C), or with venetoclax resistance (Figure 7E).

We also tested our combination treatment in a PDX model generated by transplantation of NSG mice with an AML sample from a patient who had relapsed after venetoclax treatment (Patient #7). Upon engraftment, we treated the mice for four weeks and then monitored them for survival (Supplementary Figure S13C). Importantly, we observed a significant survival advantage in the PDX mice treated with our combination treatment compared to the untreated mice (Figure 7F), but not with single agent treatment with copanlisib or valemetostat (Supplementary Figure S13D). Furthermore, we found no significant differences in body weight between treatment groups, indicating minimal toxicity of the combination treatment (Supplementary Figure S13E,F).

To test the effects of this drug combination on healthy cells and to determine a therapeutic window for this combination treatment, we performed colony assays on healthy donor bone marrow CD34+ cells with copanlisib and valemetostat. We observed a statistically significant reduction in total colony numbers with combination treatment at the highest drug doses (Figure 7G). However, the degree of inhibition was lower for healthy CD34 cells than for most MDS or AML patient samples at the same drug doses, and all myeloid colony subtypes were still present at every dose combination (Figure 7H). This suggests that AML or high-risk MDS cells are more sensitive to this drug combination than healthy human bone marrow CD34+ cells.

## Discussion

Determining more universal therapeutics that can target LSCs and finding ways to leverage the acquired resistance mechanisms to these therapeutics are critical to improve outcomes for AML patients. The PI3K pathway is activated in a variety of cancer types, including AML, and this pathway has been targeted in other forms of cancer (25). Here we report that LSCs in AML have a dependency on PI3Kα. While our data in mouse AML models and in patient samples suggest that PI3K inactivation can inhibit AML cell proliferation, our results suggest that single agent PI3K inhibitor activity is likely to be transient, with adaptive resistance developing over several days. This could be one of the reasons why prior clinical trials testing PI3K inhibitors alone had limited efficacy (18, 49). However, combining multiple classes of inhibitors can allow for improved efficacy compared to therapies targeting a single mechanism(50).

Here we show that inactivation of PI3Kα is sufficient to inhibit AKT activity in AML cells, and this leads to myeloid differentiation and loss of LSC self-renewal. Interestingly, it has been reported that forced overexpression of a myristoylated version of AKT can also result in differentiation of AML cells via FOXO inhibition(51). Rather than contradicting our findings, these observations support the concept that tight regulation of the PI3K/AKT pathway is critical to maintain normal homeostasis, including in LSCs. Together, these data also suggest that either insufficient or excessive PI3K/AKT pathway activation can impair LSC maintenance.

We also demonstrate that leukemic cells can develop adaptive resistance to PI3K inhibition via a non-genetic mechanism, whereby EZH2 protein levels are downregulated over time in response to PI3K/AKT inactivation. In murine models of KMT2A-MLLT3 AML, inactivation of EZH2 and other components of PRC2 compromises leukemic proliferation and prolongs survival (45, 52). However, Gollner and colleagues previously reported that proteasomal degradation of EZH2 can be a mechanism of chemotherapy resistance in AML (53). Our results suggest that loss of EZH2 protein can also give the leukemic cells a survival advantage in the setting of PI3K inactivation. Interestingly, a recent report also identified a similar resistance mechanism with EZH2 downregulation in response to FLT3 inhibitors, but this was limited to FLT3-mutated AML(54).

Our findings add additional context to the contradictory roles of EZH2 in myeloid malignancies. As Basheer and colleagues demonstrated, the role of EZH2 is very context-dependent, with it being either a tumor suppressor or an oncogene depending on the stage of disease and depending on which of its targets are being manipulated by its activity(27). Somewhat counterintuitively, Fujita et al noted an increase in EZH2 expression in the LSC pool, whereas Basheer et al indicate that EZH2 acts as a tumor suppressor at disease onset(44). We posit that part of what dictates this context is the phosphorylation status of EZH2 at Ser21, since EHZ2 can be phosphorylated by AKT at this site (55). Given these complexities, it follows that the targeting of EZH2 and EZH1/2 in preclinical studies has had mixed results(45, 56).

Notably, Fujita and colleagues demonstrated that the genetic ablation of both EZH1 and EZH2 is required to successfully target leukemic cells and deplete the LSC pool, compared to targeting EZH2 alone(44). Our results are consistent with this requirement for dual EZH1/2 disruption in LSCs. EZH1/2 inhibitors as solo agents have had varying degrees of success in preclinical studies in AML(45, 56). Our work identifies a mechanistic basis for enhancing the efficacy of both PI3K inhibitors and EZH1/2 inhibitors in AML. We also show that we can leverage this resistance mechanism by combining a PI3K inhibitor with an EZH1/2 dual inhibitor, exploiting the new dependency of leukemic cells on EZH1 in the context of PI3K inhibition. This combination is particularly effective at depleting functional LSCs, as we observed in the NPM1c-NRAS^G12D^ and KMT2A-MLLT3 mouse models of AML.

Several PI3K inhibitors, including copanlisib (Bayer), were approved by the FDA for lymphoma, though copanlisib was recently removed from the U.S. market due to insufficient efficacy in follicular lymphoma, despite low toxicity in patients(57). Although no PI3K inhibitors are currently approved for AML treatment, several PI3K/AKT pathway inhibitors, including alpelisib and inavolisib, are approved for other malignancies and have reasonable toxicity profiles in patients (58, 59). The EZH1/2 inhibitor valemetostat (Daichi Sankyo) is approved in Japan for the treatment of adult T cell leukemia-lymphoma (ATLL), and was also well-tolerated in patients (57, 60). While the specific combination of a PI3K inhibitor and valemetostat has not yet been tested in patients, our colony assays on mouse bone marrow cells and healthy donor CD34 cells, and combination drug treatment in multiple mouse AML models suggest that a therapeutic window for this combination treatment should be achievable.

In the last decade, there have been advancements in the targeting of epigenetic mechanisms in AML therapies, typically in combination with other therapeutics(3). By simultaneously targeting an oncogenic signaling pathway and an epigenetic regulator, our approach enables more effective targeting across diverse AML subtypes. Our work suggests that a broad range of AML cases may be susceptible to the combination of a PI3K inhibitor and an EZH1/2 inhibitor. However, more studies will be needed to elucidate the specific determinants of response to this drug combination in AML, and to understand which specific molecular or cytogenetic profiles could confer sensitivity to our proposed combination treatment. Importantly, our preclinical studies suggest that several difficult to treat subtypes of AML, including AML with complex cytogenetics, TP53 mutated AML, and venetoclax-resistant AML are still sensitive to the copanlisib and valemetostat combination. In summary, our data supports the combined PI3K and EZH1/2 inhibition as a promising therapeutic regimen for AML patients, especially for targeting leukemia-initiating cells, which could lead to a reduced risk of relapse.

## Acknowledgements

We thank the MDS/AML patients for their contributions. We thank Daqian Sun of the AECOM Stem Cell Isolation and Xenotransplantation Facility (funded through New York Stem Cell Science grant no. C029154) and Jinhang Zhang, M.Liu and A. Feng from the AECOM Flow Cytometry Core Facility for assistance with flow cytometry. We thank J. Grimm for assistance with generating the ELDA graph. We thank L. Gurska for critical reading of the manuscript. We thank all past and present members of the Gritsman lab for valuable discussions about this work.

## Funding

This work was supported by National Institutes of Health grants R01CA196973 (to KG) and K08CA149208 (to KG), the V Foundation V Scholar Award (to KG), the Alexandrine and Alexander L. Sinsheimer Scholar Award (to KG), startup funds from Montefiore Einstein Comprehensive Cancer Center (MECCC) (to KG); the Paul S. Frenette Scholar Awards Program of the Ruth L. and David S. Gottesman Institute for Stem Cell Research and Regenerative Medicine (to S.G.G-S.); The National Center for Advancing Translational Sciences, and the National Institutes of Health, through CTSA award number TL1TR002557 (to S.G.G-S). S.H. was partially supported by The Einstein Training Program in Stem Cell Research from the Empire State Stem Cell Fund through New York State Department of Health Contract C30292GG, as well as by Montefiore Einstein Comprehensive Cancer Center Support Grant of the NIH under award number P30CA013330. K.A. was partially supported by the NHLBI/NIH Ruth L. Kirschstein National Research Service Award F32HL146119 and the IRACDA/BETTR training Institutional Research and Academic Career Development Award 2K12GH102779-07A1. S. S. gratefully acknowledges for funding the Hevolution Foundation (AFAR), the Einstein-Mount Sinai Diabetes center, and the NIH Office of the Director (S10OD030286). This work was supported by NIH grant R35CA253127 (to U.S.). The content is solely the responsibility of the authors and does not necessarily represent the official views of the NIH. U.S. holds the Edward P. Evans Endowed Professorship in Myelodysplastic Syndromes at Albert Einstein College of Medicine. The Endowed Professorship was supported by a grant from the Edward P. Evans Foundation. For flow cytometry, this work used the analyzers Cytek Aurora Multiparameter Flow Cytometer and BD LSR-II with the help from J. Zhang, A. Fnu, and M. Liu. The Cytek Aurora Multiparameter Flow Cytometer was purchased with funding from the NIH SIG grant #1S10OD026833-01.

## Author Contributions

S.G.G-S and K.G. planned and performed experiments, acquired funding, analyzed data, and wrote the manuscript. I.K., T.S., and S.H. planned and performed experiments and analyzed data. E.A., A.B, K.A., R.H., and M.S. performed experiments. S.S. assisted with experimental design and data interpretation. S-R. N. performed experiments. J.V., S.K., U.S., A.V., S.C., L.M., and A.S. provided materials and assisted with data interpretation. All authors read and approved the manuscript.

## Competing Interests

K.G. received research funding from iOnctura and ADC Therapeutics. U.S. has received research funding from GlaxoSmithKline, Bayer Healthcare, Aileron Therapeutics, and Novartis; has received compensation for consultancy services and for serving on scientific advisory boards from GlaxoSmithKline, Bayer Healthcare, Celgene, Aileron Therapeutics, Novartis, Stelexis Therapeutics, Pieris Pharmaceuticals, and Trillium Therapeutics; and has equity ownership in and is serving on the board of directors of Stelexis Therapeutics; all outside of the reported work. A.S. has received research funding from Kymera Therapeutics, advisory board fees from Gilead Sciences, Rigel Pharmaceuticals and Kymera Therapeutics, consultancy fees from Janssen Pharmaceuticals and honoraria from National Association of Continuing Education & PeerView. A.V. has received research funding from GlaxoSmithKline, BMS, Jannsen, Incyte, MedPacto, Celgene, Novartis, Curis, Prelude and Eli Lilly and Company, has received compensation as a scientific advisor to s, Stelexis Therapeutics, Calico, Acceleron Pharma, Aurigene and Celgene, and has equity ownership in Bioconvergent health, Throws Exception and Stelexis Therapeutics.

## Notes

https://www.ncbi.nlm.nih.gov/geo/query/acc.cgi?acc=GSE261355

### Competing Interest Statement

K.G. received research funding from iOnctura and ADC Therapeutics. U.S. has received research fund-ing from GlaxoSmithKline, Bayer Healthcare, Aileron Therapeutics, and Novartis; has received compensa-tion for consultancy services and for serving on scien-tific advisory boards from GlaxoSmithKline, Bayer Healthcare, Celgene, Aileron Therapeutics, Novartis, Stelexis Therapeutics, Pieris Pharmaceuticals, and Trillium Therapeutics; and has equity ownership in and is serving on the board of directors of Stelexis Therapeutics; all outside of the reported work. A.S. has received research funding from Kymera Thera-peutics, advisory board fees from Gilead Sciences, Rigel Pharmaceuticals and Kymera Therapeutics, con-sultancy fees from Janssen Pharmaceuticals and hon-oraria from National Association of Continuing Edu-cation & PeerView. A.V. has received research fund-ing from GlaxoSmithKline, BMS, Jannsen, Incyte, MedPacto, Celgene, Novartis, Curis, Prelude and Eli Lilly and Company, has received compensation as a scientific advisor to s, Stelexis Therapeutics, Calico, Acceleron Pharma, Aurigene and Celgene, and has equity ownership in Bioconvergent health, Throws Exception and Stelexis Therapeutics. The content is solely the responsibility of the authors and does not necessarily represent the official views of the National Institutes of Health.

### Summary of Updates

Figures and text were extensively revised

## References

1. Heuser M, Ofran Y, Boissel N, Brunet Mauri S, Craddock C, Janssen J, et al. Acute myeloid leukaemia in adult patients: ESMO Clinical Practice Guidelines for diagnosis, treatment and follow-up. Ann Oncol. 2020;31(6):697–712.

2. Stelmach P, Trumpp A. Leukemic stem cells and therapy resistance in acute myeloid leukemia. Haematologica. 2023;108(2):353–66.

3. Fennell KA, Vassiliadis D, Lam EYN, Martelotto LG, Balic JJ, Hollizeck S, et al. Non-genetic determinants of malignant clonal fitness at single-cell resolution. Nature. 2022;601(7891):125–31.

4. Chen SJ, Shen Y, Chen Z. A panoramic view of acute myeloid leukemia. Nature genetics. 2013;45(6):586–7.

5. Nepstad I, Hatfield KJ, Gronningsaeter IS, Reikvam H. The PI3K-Akt-mTOR Signaling Pathway in Human Acute Myeloid Leukemia (AML) Cells. Int J Mol Sci. 2020;21(8).

6. Grandage VL, Gale RE, Linch DC, Khwaja A. PI3-kinase/Akt is constitutively active in primary acute myeloid leukaemia cells and regulates survival and chemoresistance via NF-kappaB, Mapkinase and p53 pathways. Leukemia. 2005;19(4):586–94.

7. Gurska LM, Ames K, Gritsman K. Signaling Pathways in Leukemic Stem Cells. Adv Exp Med Biol. 2019;1143:1–39.

8. Min YH, Eom JI, Cheong JW, Maeng HO, Kim JY, Jeung HK, et al. Constitutive phosphorylation of Akt/PKB protein in acute myeloid leukemia: its significance as a prognostic variable. Leukemia. 2003;17(5):995–7.

9. Kharas MG, Okabe R, Ganis JJ, Gozo M, Khandan T, Paktinat M, et al. Constitutively active AKT depletes hematopoietic stem cells and induces leukemia in mice. Blood. 2010;115(7):1406–15.

10. Yilmaz OH, Valdez R, Theisen BK, Guo W, Ferguson DO, Wu H, et al. Pten dependence distinguishes haematopoietic stem cells from leukaemia-initiating cells. Nature. 2006;441(7092):475–82.

11. Zhang J, Grindley JC, Yin T, Jayasinghe S, He XC, Ross JT, et al. PTEN maintains haematopoietic stem cells and acts in lineage choice and leukaemia prevention. Nature. 2006;441(7092):518–22.

12. Bilanges B, Posor Y, Vanhaesebroeck B. PI3K isoforms in cell signalling and vesicle trafficking. Nat Rev Mol Cell Biol. 2019;20(9):515–34.

13. Vanhaesebroeck B, Whitehead MA, Pineiro R. Molecules in medicine mini-review: isoforms of PI3K in biology and disease. J Mol Med (Berl). 2016;94(1):5–11.

14. Lu JW, Lin YM, Lai YL, Chen CY, Hu CY, Tien HF, et al. MK-2206 induces apoptosis of AML cells and enhances the cytotoxicity of cytarabine. Med Oncol. 2015;32(7):206.

15. Park S, Chapuis N, Bardet V, Tamburini J, Gallay N, Willems L, et al. PI-103, a dual inhibitor of Class IA phosphatidylinositide 3-kinase and mTOR, has antileukemic activity in AML. Leukemia. 2008;22(9):1698–706.

16. Jin L, Tabe Y, Kojima K, Shikami M, Benito J, Ruvolo V, et al. PI3K inhibitor GDC-0941 enhances apoptotic effects of BH-3 mimetic ABT-737 in AML cells in the hypoxic bone marrow microenvironment. J Mol Med (Berl). 2013;91(12):1383–97.

17. Sandhofer N, Metzeler KH, Rothenberg M, Herold T, Tiedt S, Groiss V, et al. Dual PI3K/mTOR inhibition shows antileukemic activity in MLL-rearranged acute myeloid leukemia. Leukemia. 2015;29(4):828–38.

18. Fransecky L, Mochmann LH, Baldus CD. Outlook on PI3K/AKT/mTOR inhibition in acute leukemia. Mol Cell Ther. 2015;3:2.

19. Hemmati S, Sinclair T, Tong M, Bartholdy B, Okabe RO, Ames K, et al. PI3 kinase alpha and delta promote hematopoietic stem cell activation. JCI Insight. 2019;5(13).

20. Yuzugullu H, Baitsch L, Von T, Steiner A, Tong H, Ni J, et al. A PI3K p110beta-Rac signalling loop mediates Pten-loss-induced perturbation of haematopoiesis and leukaemogenesis. Nat Commun. 2015;6:8501.

21. Gritsman K, Yuzugullu H, Von T, Yan H, Clayton L, Fritsch C, et al. Hematopoiesis and RAS-driven myeloid leukemia differentially require PI3K isoform p110alpha. J Clin Invest. 2014;124(4):1794–809.

22. Luo Q, Raulston EG, Prado MA, Wu X, Gritsman K, Whalen KS, et al. Targetable leukaemia dependency on noncanonical PI3Kgamma signalling. Nature. 2024;630(8015):198–205.

23. Gu H, Chen C, Hou ZS, He XD, Xie S, Ni J, et al. PI3Kgamma maintains the self-renewal of acute myeloid leukemia stem cells by regulating the pentose phosphate pathway. Blood. 2024;143(19):1965–79.

24. Kelly LM, Rutter JC, Lin KH, Ling F, Duchmann M, Latour E, et al. Targeting a lineage-specific PI3K -Akt signaling module in acute myeloid leukemia using a heterobifunctional degrader molecule. Nat Cancer. 2024;5(7):1082–101.

25. Vanhaesebroeck B, Perry MWD, Brown JR, Andre F, Okkenhaug K. PI3K inhibitors are finally coming of age. Nat Rev Drug Discov. 2021;20(10):741–69.

26. Rinke J, Chase A, Cross NCP, Hochhaus A, Ernst T. EZH2 in Myeloid Malignancies. Cells. 2020;9(7).

27. Basheer F, Giotopoulos G, Meduri E, Yun H, Mazan M, Sasca D, et al. Contrasting requirements during disease evolution identify EZH2 as a therapeutic target in AML. J Exp Med. 2019;216(4):966–81.

28. Mochizuki-Kashio M, Aoyama K, Sashida G, Oshima M, Tomioka T, Muto T, et al. Ezh2 loss in hematopoietic stem cells predisposes mice to develop heterogeneous malignancies in an Ezh1-dependent manner. Blood. 2015;126(10):1172–83.

29. Gu Z, Liu Y, Cai F, Patrick M, Zmajkovic J, Cao H, et al. Loss of EZH2 Reprograms BCAA Metabolism to Drive Leukemic Transformation. Cancer Discov. 2019;9(9):1228–47.

30. Ames K, Kaur I, Shi Y, Tong MM, Sinclair T, Hemmati S, et al. PI3-kinase deletion promotes myelodysplasia by dysregulating autophagy in hematopoietic stem cells. Sci Adv. 2023;9(8):eade8222.

31. Choudhary GS, Al-Harbi S, Mazumder S, Hill BT, Smith MR, Bodo J, et al. MCL-1 and BCL-xL-dependent resistance to the BCL-2 inhibitor ABT-199 can be overcome by preventing PI3K/AKT/mTOR activation in lymphoid malignancies. Cell Death Dis. 2015;6(1):e1593.

32. Sidoli S, Bhanu NV, Karch KR, Wang X, Garcia BA. Complete Workflow for Analysis of Histone Post-translational Modifications Using Bottom-up Mass Spectrometry: From Histone Extraction to Data Analysis. J Vis Exp. 2016(111).

33. Yuan ZF, Sidoli S, Marchione DM, Simithy J, Janssen KA, Szurgot MR, et al. EpiProfile 2.0: A Computational Platform for Processing Epi-Proteomics Mass Spectrometry Data. J Proteome Res. 2018;17(7):2533–41.

34. Krivtsov AV, Armstrong SA. MLL translocations, histone modifications and leukaemia stem-cell development. Nature reviews Cancer. 2007;7(11):823–33.

35. Furet P, Guagnano V, Fairhurst RA, Imbach-Weese P, Bruce I, Knapp M, et al. Discovery of NVP-BYL719 a potent and selective phosphatidylinositol-3 kinase alpha inhibitor selected for clinical evaluation. Bioorg Med Chem Lett. 2013;23(13):3741–8.

36. Chaussade C, Rewcastle GW, Kendall JD, Denny WA, Cho K, Gronning LM, et al. Evidence for functional redundancy of class IA PI3K isoforms in insulin signalling. Biochem J. 2007;404(3):449–58.

37. Herman SE, Gordon AL, Wagner AJ, Heerema NA, Zhao W, Flynn JM, et al. Phosphatidylinositol 3-kinase-delta inhibitor CAL-101 shows promising preclinical activity in chronic lymphocytic leukemia by antagonizing intrinsic and extrinsic cellular survival signals. Blood. 2010;116(12):2078–88.

38. Liu N, Rowley BR, Bull CO, Schneider C, Haegebarth A, Schatz CA, et al. BAY 80-6946 is a highly selective intravenous PI3K inhibitor with potent p110alpha and p110delta activities in tumor cell lines and xenograft models. Mol Cancer Ther. 2013;12(11):2319–30.

39. Maira SM, Pecchi S, Huang A, Burger M, Knapp M, Sterker D, et al. Identification and characterization of NVP-BKM120, an orally available pan-class I PI3-kinase inhibitor. Mol Cancer Ther. 2012;11(2):317–28.

40. Jia S, Liu Z, Zhang S, Liu P, Zhang L, Lee SH, et al. Essential roles of PI(3)K-p110beta in cell growth, metabolism and tumorigenesis. Nature. 2008;454(7205):776–9.

41. Krivtsov AV, Twomey D, Feng Z, Stubbs MC, Wang Y, Faber J, et al. Transformation from committed progenitor to leukaemia stem cell initiated by MLL-AF9. Nature. 2006;442(7104):818–22.

42. Subramanian A, Tamayo P, Mootha VK, Mukherjee S, Ebert BL, Gillette MA, et al. Gene set enrichment analysis: a knowledge-based approach for interpreting genome-wide expression profiles. Proc Natl Acad Sci U S A. 2005;102(43):15545–50.

43. Castanza AS, Recla JM, Eby D, Thorvaldsdottir H, Bult CJ, Mesirov JP. Extending support for mouse data in the Molecular Signatures Database (MSigDB). Nat Methods. 2023;20(11):1619–20.

44. Fujita S, Honma D, Adachi N, Araki K, Takamatsu E, Katsumoto T, et al. Dual inhibition of EZH1/2 breaks the quiescence of leukemia stem cells in acute myeloid leukemia. Leukemia. 2018;32(4):855–64.

45. Xu B, On DM, Ma A, Parton T, Konze KD, Pattenden SG, et al. Selective inhibition of EZH2 and EZH1 enzymatic activity by a small molecule suppresses MLL-rearranged leukemia. Blood. 2015;125(2):346–57.

46. Chakraborty S, Morganti C, Zaldana K, Rivera Pena B, Zhang H, Verma D, et al. A STAT3 degrader demonstrates efficacy in venetoclax resistant acute myeloid leukemia. Leukemia. 2026.

47. Dovey OM, Cooper JL, Mupo A, Grove CS, Lynn C, Conte N, et al. Molecular synergy underlies the co-occurrence patterns and phenotype of NPM1-mutant acute myeloid leukemia. Blood. 2017;130(17):1911–22.

48. Hu Y, Smyth GK. ELDA: extreme limiting dilution analysis for comparing depleted and enriched populations in stem cell and other assays. J Immunol Methods. 2009;347(1-2):70–8.

49. Ragon BK, Kantarjian H, Jabbour E, Ravandi F, Cortes J, Borthakur G, et al. Buparlisib, a PI3K inhibitor, demonstrates acceptable tolerability and preliminary activity in a phase I trial of patients with advanced leukemias. Am J Hematol. 2017;92(1):7–11.

50. Bayat Mokhtari R, Homayouni TS, Baluch N, Morgatskaya E, Kumar S, Das B, et al. Combination therapy in combating cancer. Oncotarget. 2017;8(23):38022–43.

51. Sykes SM, Lane SW, Bullinger L, Kalaitzidis D, Yusuf R, Saez B, et al. AKT/FOXO signaling enforces reversible differentiation blockade in myeloid leukemias. Cell. 2011;146(5):697–708.

52. Neff T, Sinha AU, Kluk MJ, Zhu N, Khattab MH, Stein L, et al. Polycomb repressive complex 2 is required for MLL-AF9 leukemia. Proc Natl Acad Sci U S A. 2012;109(13):5028–33.

53. Gollner S, Oellerich T, Agrawal-Singh S, Schenk T, Klein HU, Rohde C, et al. Loss of the histone methyltransferase EZH2 induces resistance to multiple drugs in acute myeloid leukemia. Nat Med. 2017;23(1):69–78.

54. Sung PJ, Selvam M, Riedel SS, Xie HM, Bryant K, Manning B, et al. FLT3 tyrosine kinase inhibition modulates PRC2 and promotes differentiation in acute myeloid leukemia. Leukemia. 2024;38(2):291–301.

55. Cha TL, Zhou BP, Xia W, Wu Y, Yang CC, Chen CT, et al. Akt-mediated phosphorylation of EZH2 suppresses methylation of lysine 27 in histone H3. Science. 2005;310(5746):306–10.

56. Porazzi P, Petruk S, Pagliaroli L, De Dominici M, Deming D, 2nd, Puccetti MV, et al. Targeting Chemotherapy to Decondensed H3K27me3-Marked Chromatin of AML Cells Enhances Leukemia Suppression. Cancer Res. 2022;82(3):458–71.

57. Cheson BD, O’Brien S, Ewer MS, Goncalves MD, Farooki A, Lenz G, et al. Optimal Management of Adverse Events From Copanlisib in the Treatment of Patients With Non-Hodgkin Lymphomas. Clin Lymphoma Myeloma Leuk. 2019;19(3):135–41.

58. Sirico M, D’Angelo A, Gianni C, Casadei C, Merloni F, De Giorgi U. Current State and Future Challenges for PI3K Inhibitors in Cancer Therapy. Cancers (Basel). 2023;15(3).

59. Yu M, Chen J, Xu Z, Yang B, He Q, Luo P, et al. Development and safety of PI3K inhibitors in cancer. Arch Toxicol. 2023;97(3):635–50.

60. Izutsu K, Makita S, Nosaka K, Yoshimitsu M, Utsunomiya A, Kusumoto S, et al. An open-label, single-arm phase 2 trial of valemetostat for relapsed or refractory adult T-cell leukemia/lymphoma. Blood. 2023;141(10):1159–68.

