## Supplemental Figures for "Exploiting an Epigenetic Resistance Mechanism to PI3 Kinase Inhibition in Leukemic Stem Cells"

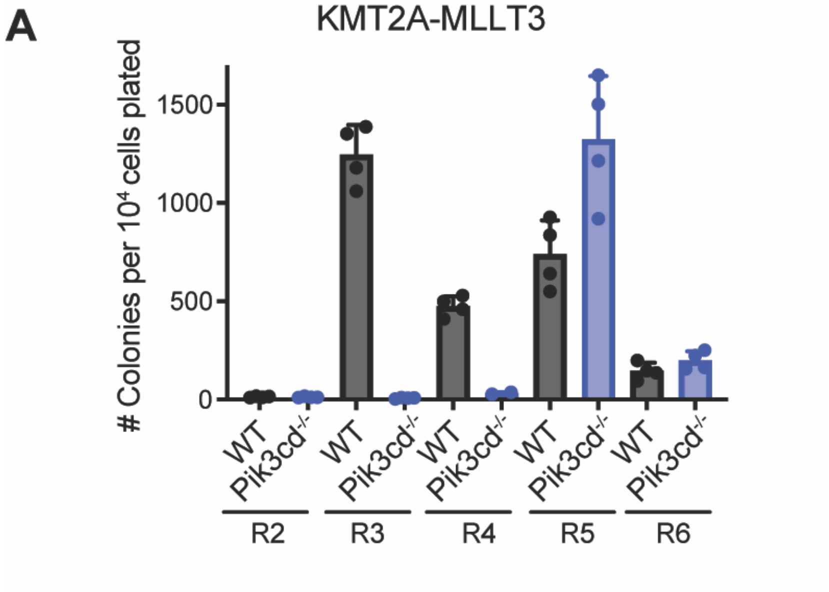


**Figure S1:** Serial replating results of KMT2A-MLLT3 cells with *Pik3cd*-/- and WT.

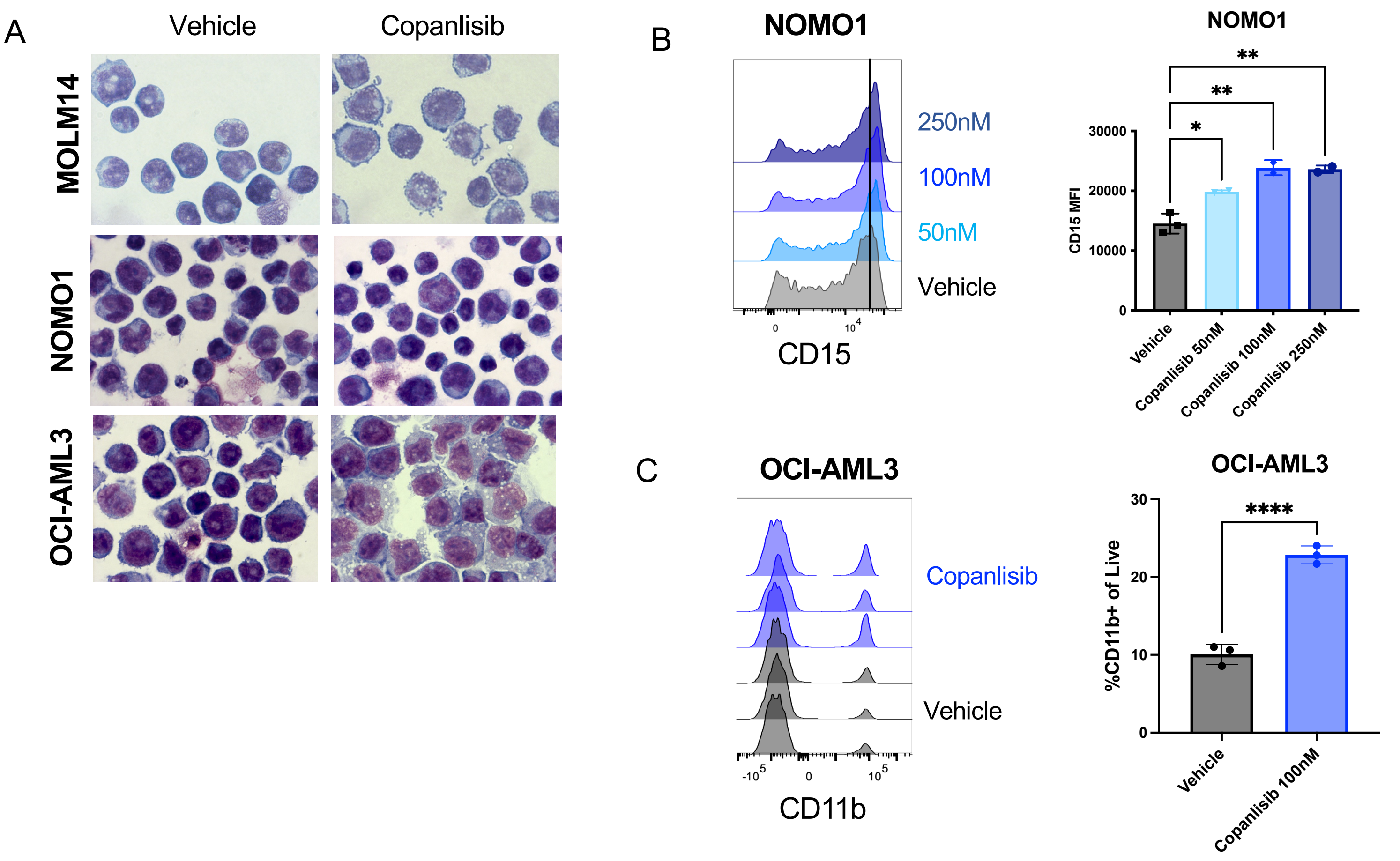


**Figure S2: PI3K Disruption Impairs Leukemic Stem Cell Self-Renewal and Promotes Myeloid Differentiation (A)** Representative images of Wright Giemsa stained cytospins from AML cell lines treated with vehicle or 100nM copanlisib for 3 days **(B)** Representative histograms of CD15 expression on NOMO1 cells treated with 100nM copanlisib for 3 days. Quantification is shown on the right. One-way ANOVA test with Tukey's multiple comparisons was used. **P<0.01 *P<0.05 **(C)** Representative flow plots and quantification of CD11b expression on OCI-AML3 cells treated with 100nM copanlisib for 3 days. Quantification is shown on the right. Unpaired t-test was used. ****P<0.0001 Each value is presented as mean +/- standard error of the mean (SEM).

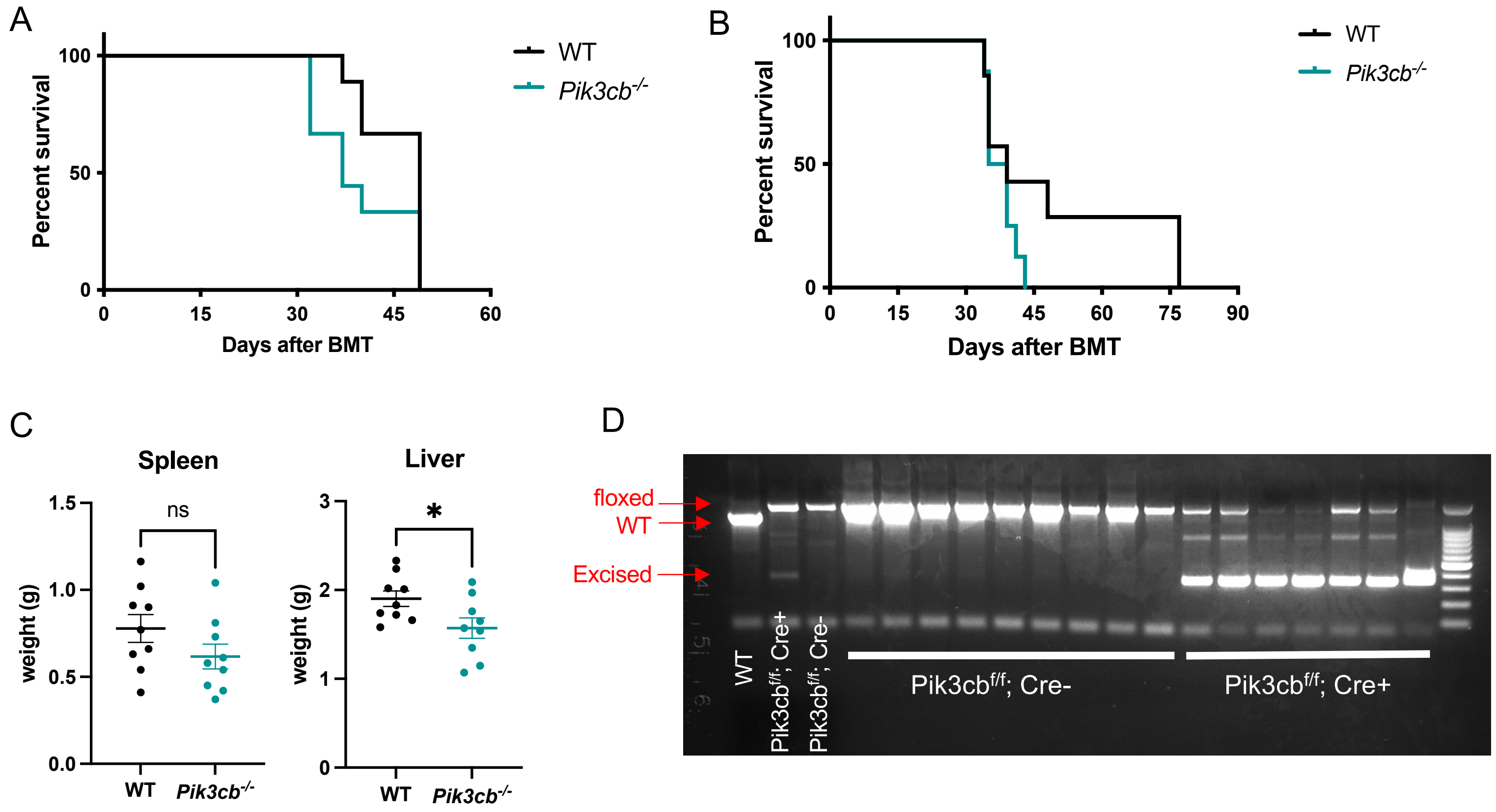


**Figure S3: p110β (*Pik3cb)* is not required for MLL-AF9 AML progression or LIC activity *in vivo***
**(A)** Kaplan-Meier survival curve of primary bone marrow transplant (BMT) mice injected with *Pik3cb*-lox-lox;Mx1-Cre or Cre- control (WT) LSK cells transduced with KMT2A-MLLT3 GFP (see Figure 2A). Log-rank analysis was used. **(B)** Kaplan-Meier survival curve of secondary BMT recipients injected with 10,000 KMT2A-MLLT3 GFP+ leukemic cells from *Pik3cb*-/- or WT primary recipient mice. Log-rank analysis was used. **(C)** Spleen and liver weights from primary BMT mice. Each value is presented as mean +/- standard error of the mean (SEM). Unpaired t-test was used. *P<0.05 **(D)** Genotyping results for primary transplant recipients post pIpC injection showing excision of *Pik3cb* in leukemic cells from Mx1-Cre+ animals

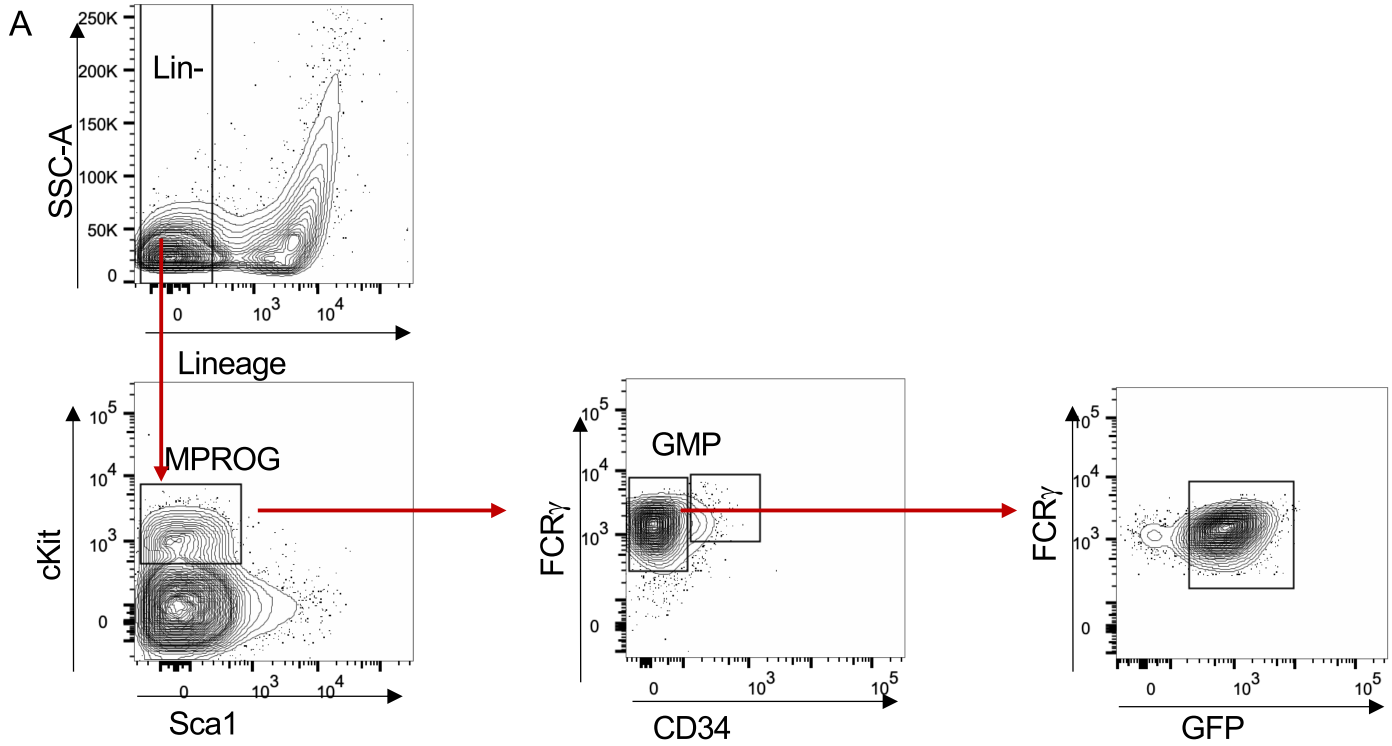


**Figure S4:** Gating strategy for LSCs in mouse bone marrow

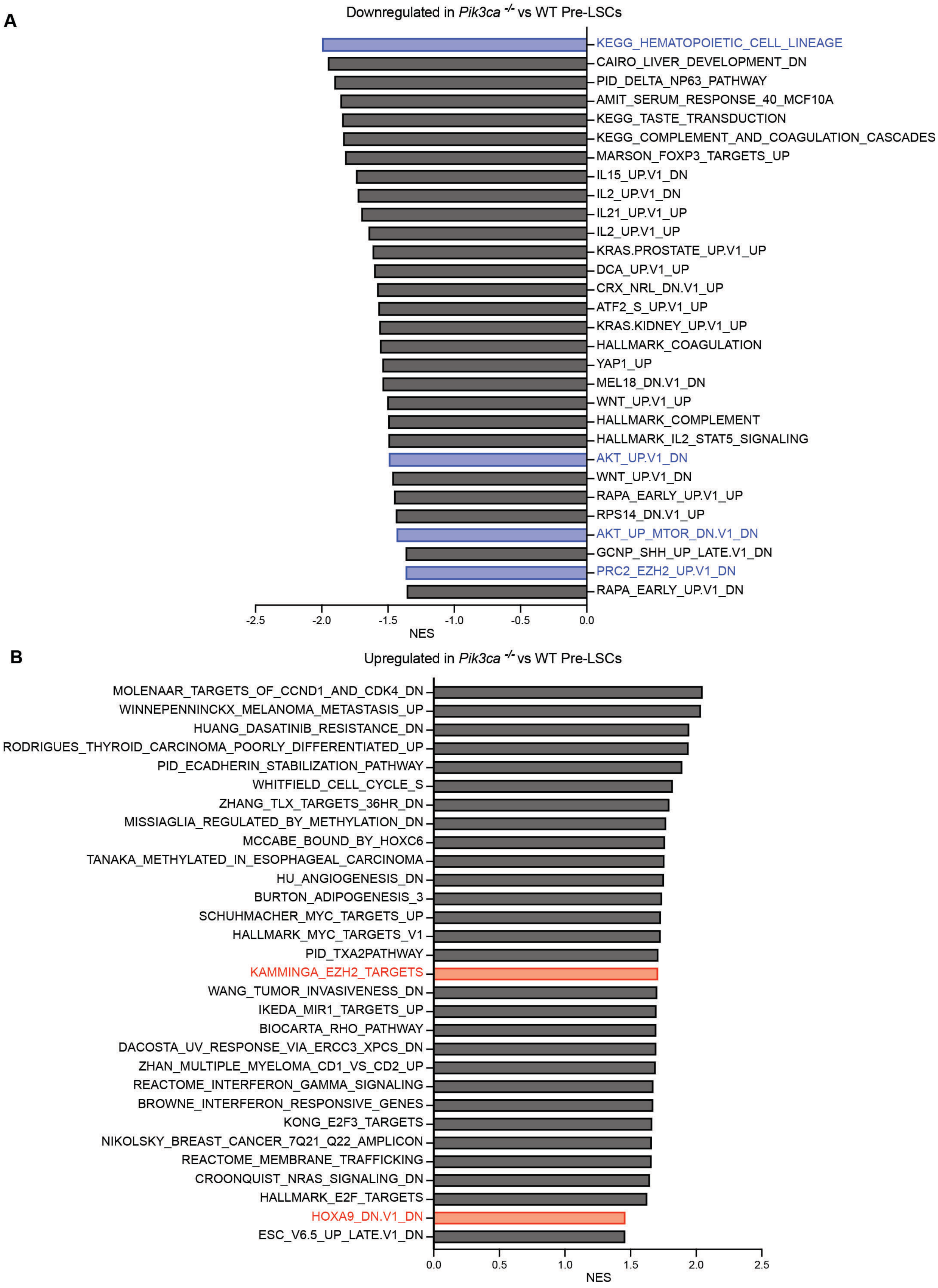


**Figure S5: *Pik3ca* Deletion Alters pre-LSC Gene Expression Signatures, Including in Hematopoietic Lineage specification and PRC2 regulation**

Gene set enrichment analysis (GSEA) was performed on the microarray dataset from preclinical LSCs (pre-LSCs: GFP+ GMPs) sorted from KMT2A-MLLT3 recipient bone marrow at 45 days post-transplantation. Cutoff of p-value <0.05 and lowest 30 FDR-q values depicted **(A)** Negatively enriched gene sets **(B)** Positively enriched gene sets

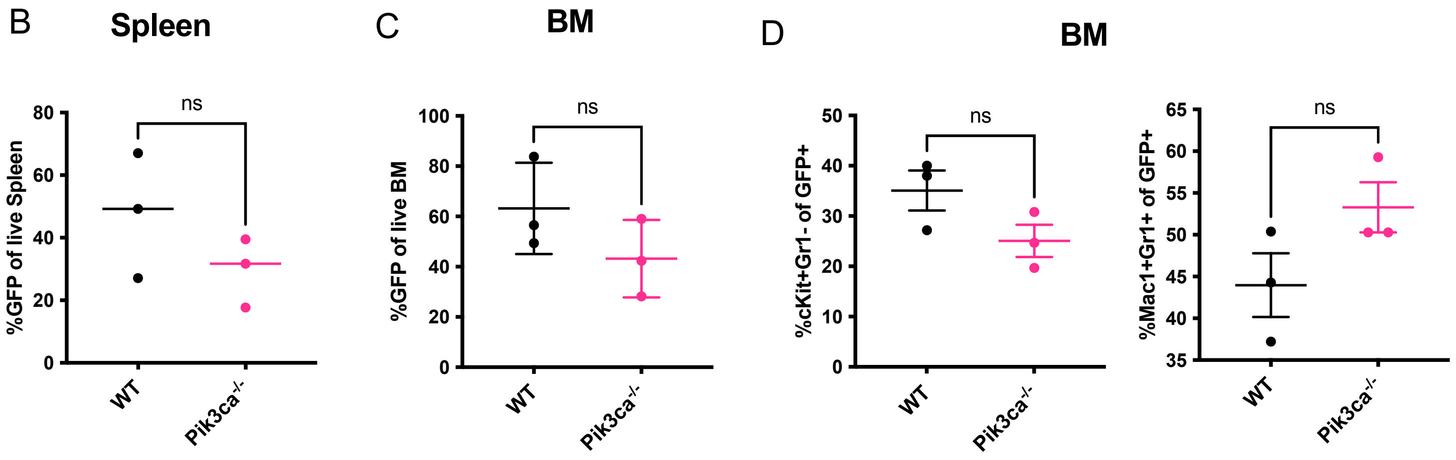


**Figure S6: There is no significant difference in the disease phenotype at time of death between WT and *Pik3ca*-/- leukemic mice** . (A) Quantification of %GFP of live spleen cells in leukemic mice **(B)** Quantification of %GFP of live bone marrow cells in leukemic mice. **(C)** Quantification of cell populations within the leukemic GFP+ compartment in the bone marrow of leukemic mice. Each value is presented as mean+/- standard error of the mean (SEM). Unpaired t-test was used to compare two groups.

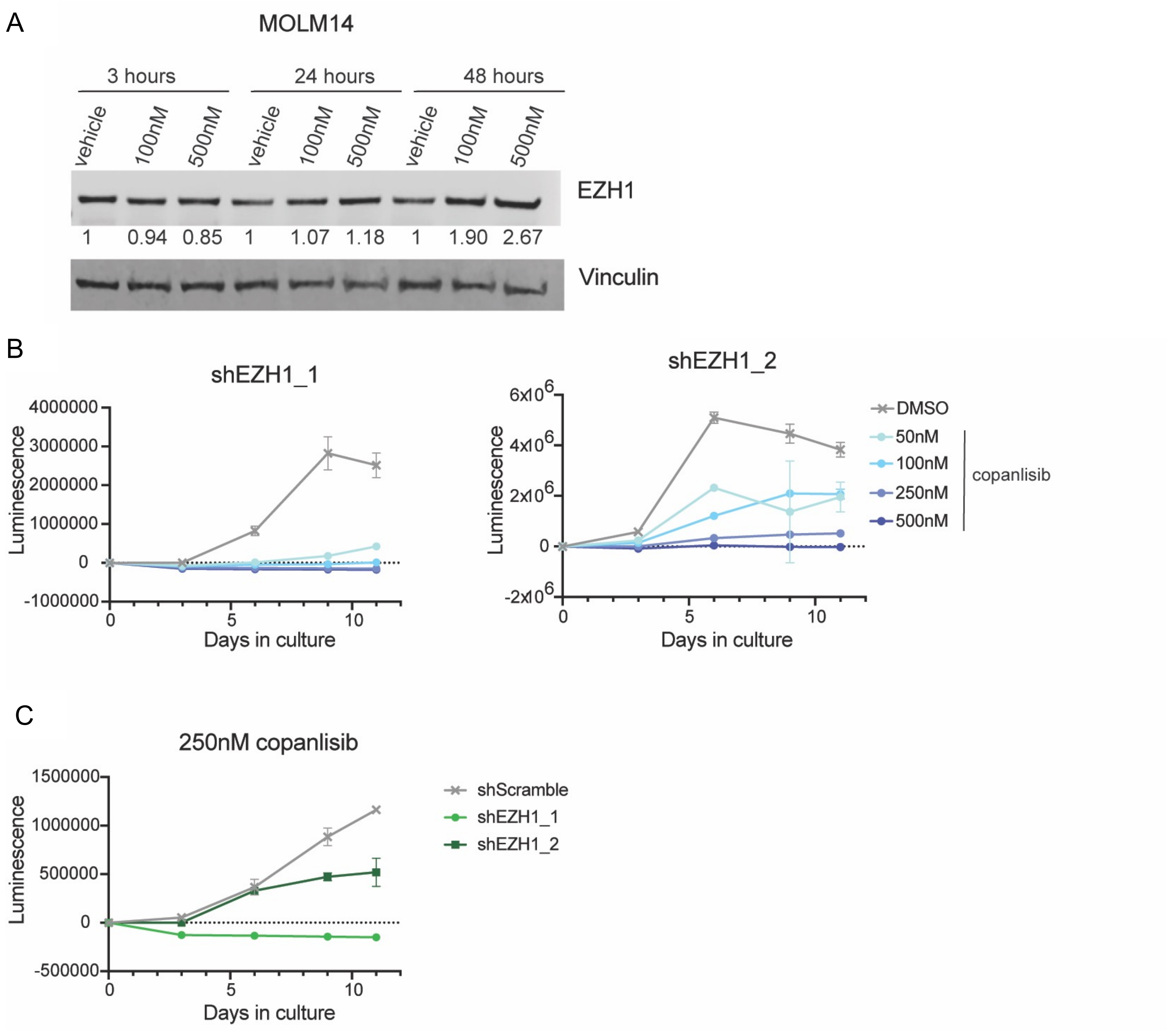


**Figure S7: EZH1 is Essential After PI3K Inhibition (A)** Western blot analysis for EZH1 in copanlisib-treaeted MOLM14 cells. Quantification is shown below each band, normalized to vehicle control over vinculin loading control. **(B)** Proliferation assays on NOMO1 cells with shEZH1 knockdown with copanlisib treatment **(C)** Proliferation assay comparing copanlisib-treated NOMO1 cells with shEZH1 knockdown.

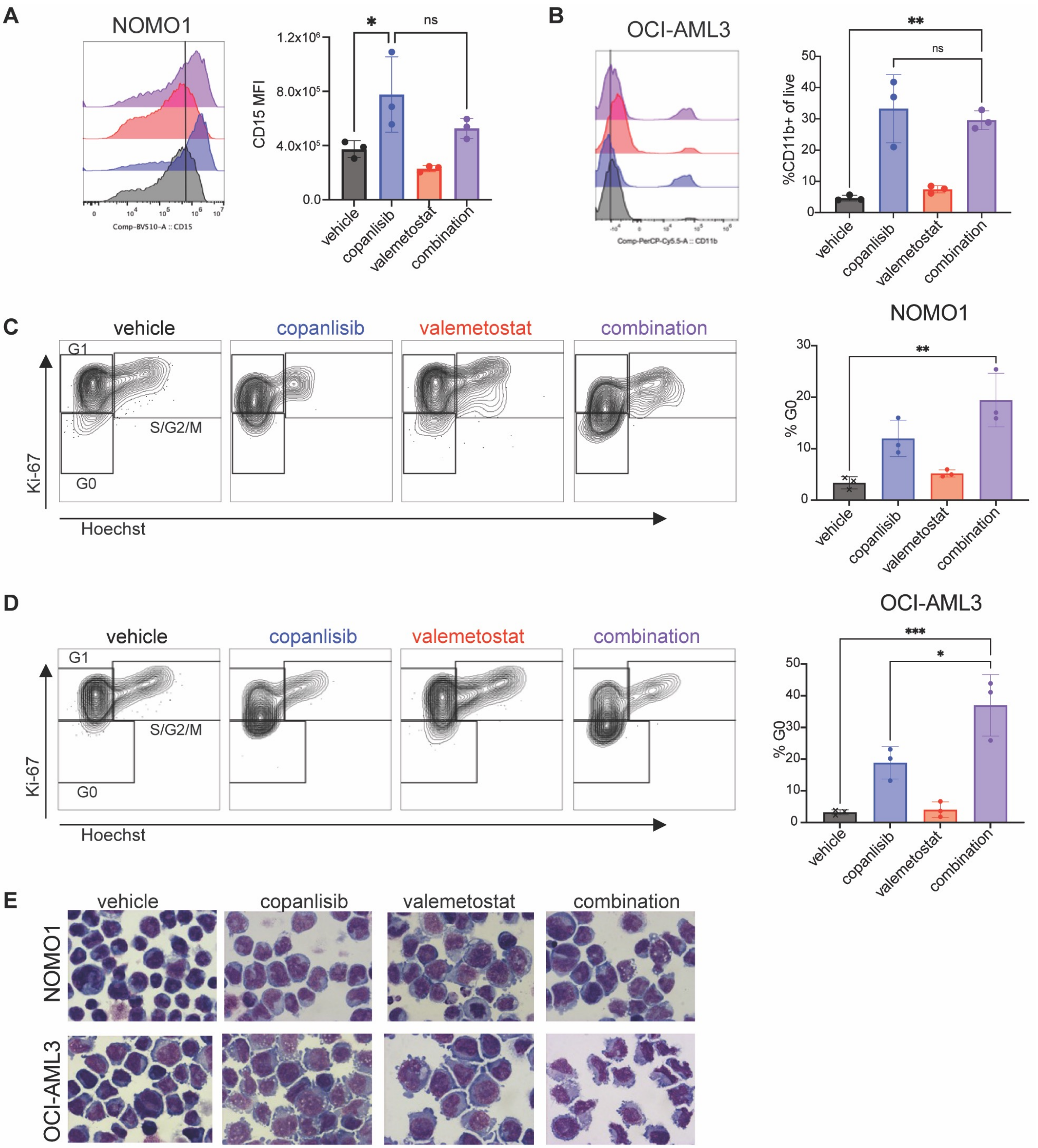


**Figure S8: PI3K Inhibition Cooperates with EZH1/2 Dual Inhibition (A-B)** Representative histograms and quantification of median fluorescence intensity (MFI) of monocytic markers CD115 **(A)** and CD11b **(B)** in two different AML cell lines after treatment with either 100nM copanlisib, 500nM valemetostat, or the combination after 6 days of treatment. **(C-D)** Representative flow cytometry plots and quantitative histograms of cell cycle analysis on NOMO1 cells **(C)** or OCI-AML3 cells **(D)** after 3 days of treatment with either 100nM copanlisib, 500nM valemetostat, or the combination. Each value is presented as mean +/- standard error of the mean (SEM). One-way ANOVA test with Tukey's multiple comparisons was used in B-D. ***P<0.001 **P<0.01 **(E)** Representative images of Wright Giemsa stained cytospins of NOMO1 or OCI-AML3 cells treated for 6 days with either 100nM copanlisib, 500nM valemetostat, or the combination.

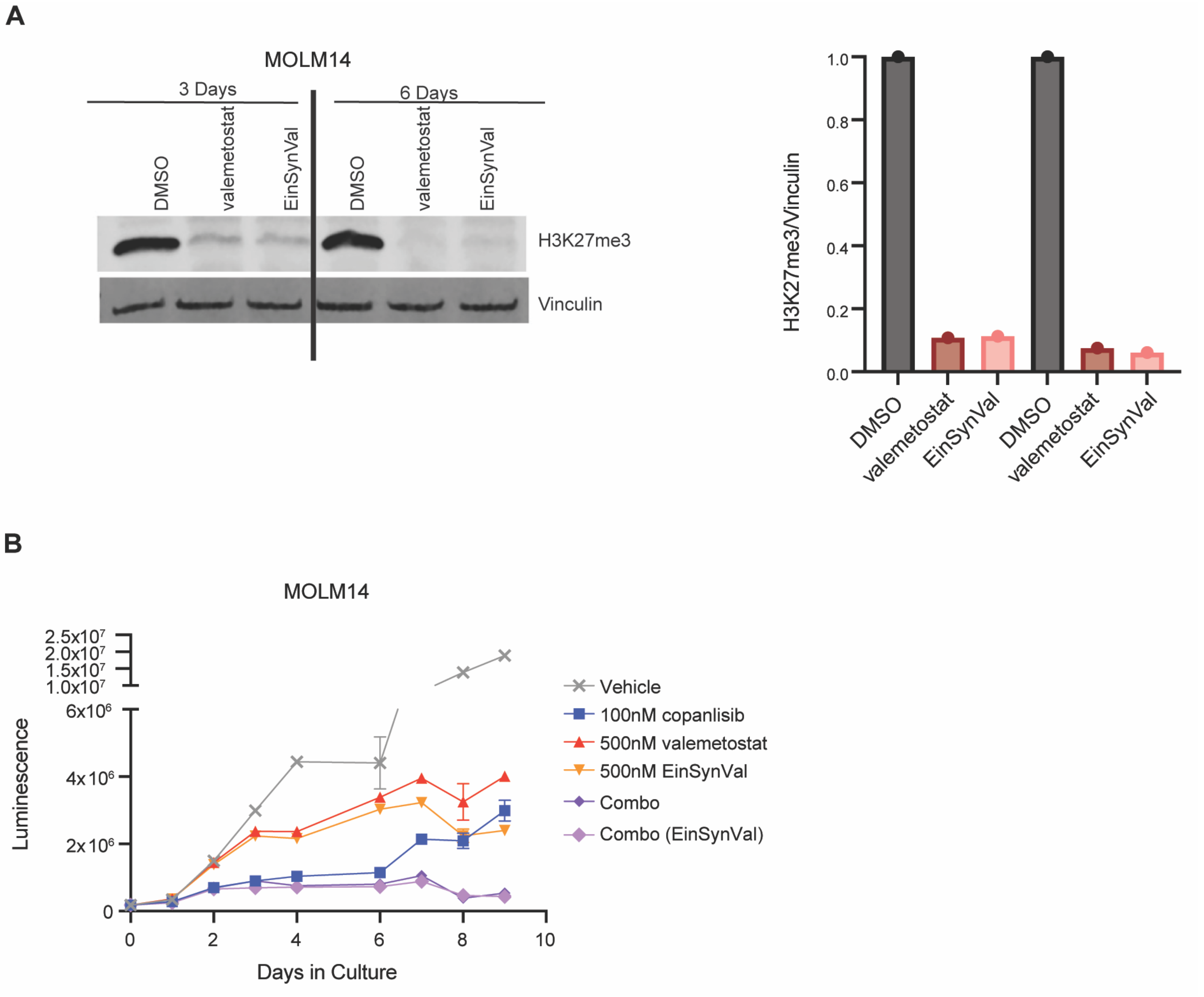


**Figure S9: Validation of Synthesized EZH1/2 Inhibitor (A)** Western blot comparing H3K27me3 levels in MOLM14 cells treated with Valemetostat (Chemietek) or DS-3201 synthesized at Albert Einstein (EinSynVal) after 3 and 6 days of treatment. Quantification of Western Blot normalized to vinculin loading control is shown on the right. **(B)** CellTiterGlo proliferation assay comparing Valemetostat (chemietek) to Einstein Synthesized DS-3201.

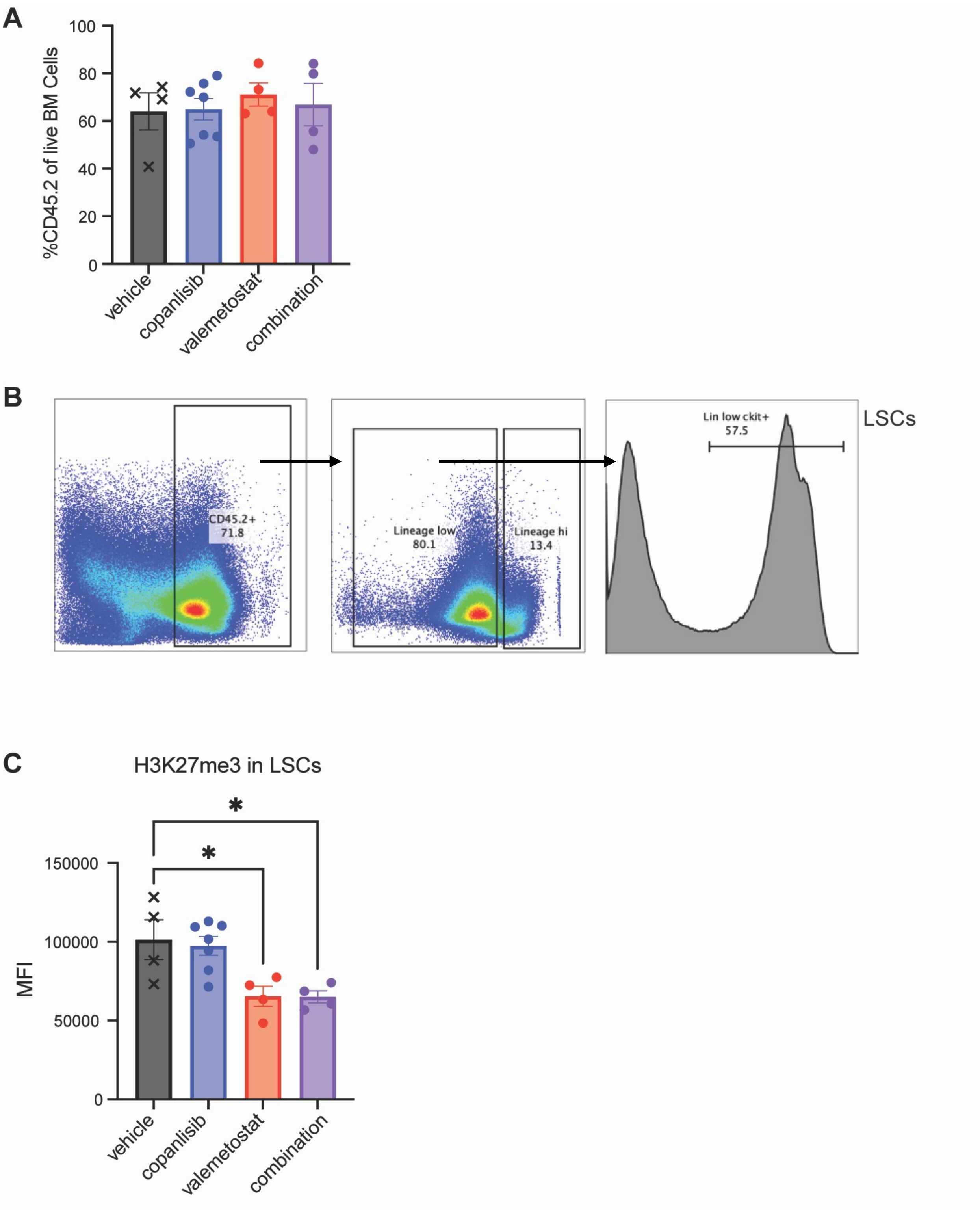


**Figure S10: EZH1/2 Inhibition Causes Impaired PRC2 Activity in LSCs (A)** Flow cytometry analysis of CD45.2 on bone marrow aspirates of NPM1c-NRAS drug treated mice after 2 weeks of treatment. **(B)** Gating strategy for LSCs in mouse bone marrow. **(C)** Quantification of Median Fluorescent Intensity (MFI) of H3K27me3 in LSCs demonstrating on target drug effects in LSCs. Each value is presented as mean +/- standard error of the mean (SEM). One-way ANOVA test with Tukey's multiple comparisons was used in C. *P≤0.05

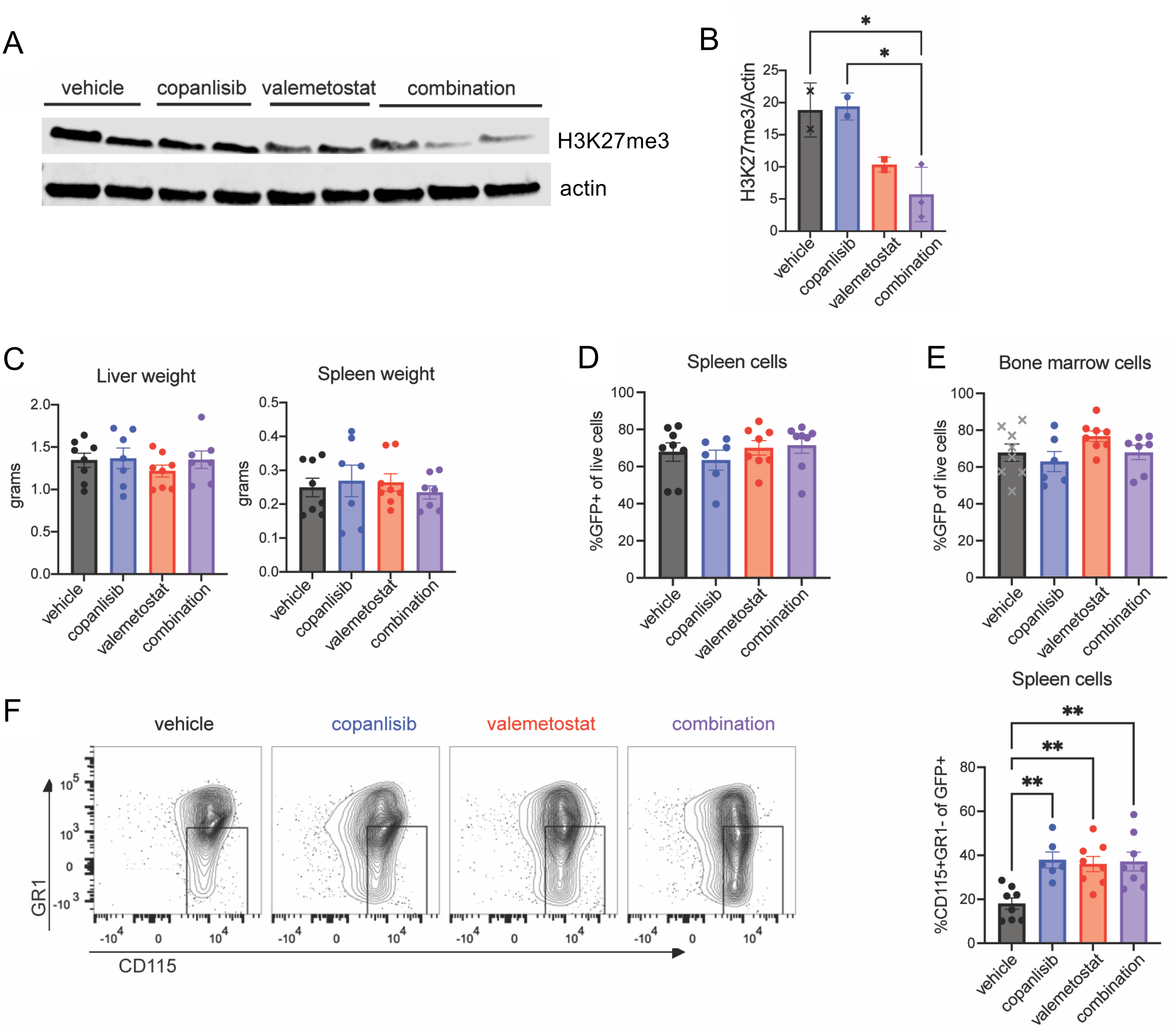


**Figure S11: PI3K Inhibition Cooperate with EZH1/2 Dual Inhibition to Target LSCs (A-C)** Western analysis on spleenocytes from primary transplant recipients (from Fig6A) demonstrating on target drug effects of valemetostat (quantified in **B**) *in vivo*. One-way ANOVA test with Tukey's multiple comparisons was used to compare signal normalized to β-actin loading control. **(C)** Organ weights measured at time of euthanasia at 2 weeks post treatment. **(D-E)** Disease burden measurement by percent of GFP+ cells in the spleen **(D)** and bone marrow **(E)** **(F)** Flow cytometry with the myeloid markers CD115 and Gr1. Representative flow plots are shown on the left. Quantification of CD115+Gr1- cells is shown on the right. $^{**}P \leq 0.01$, $^{*}P \leq 0.05$

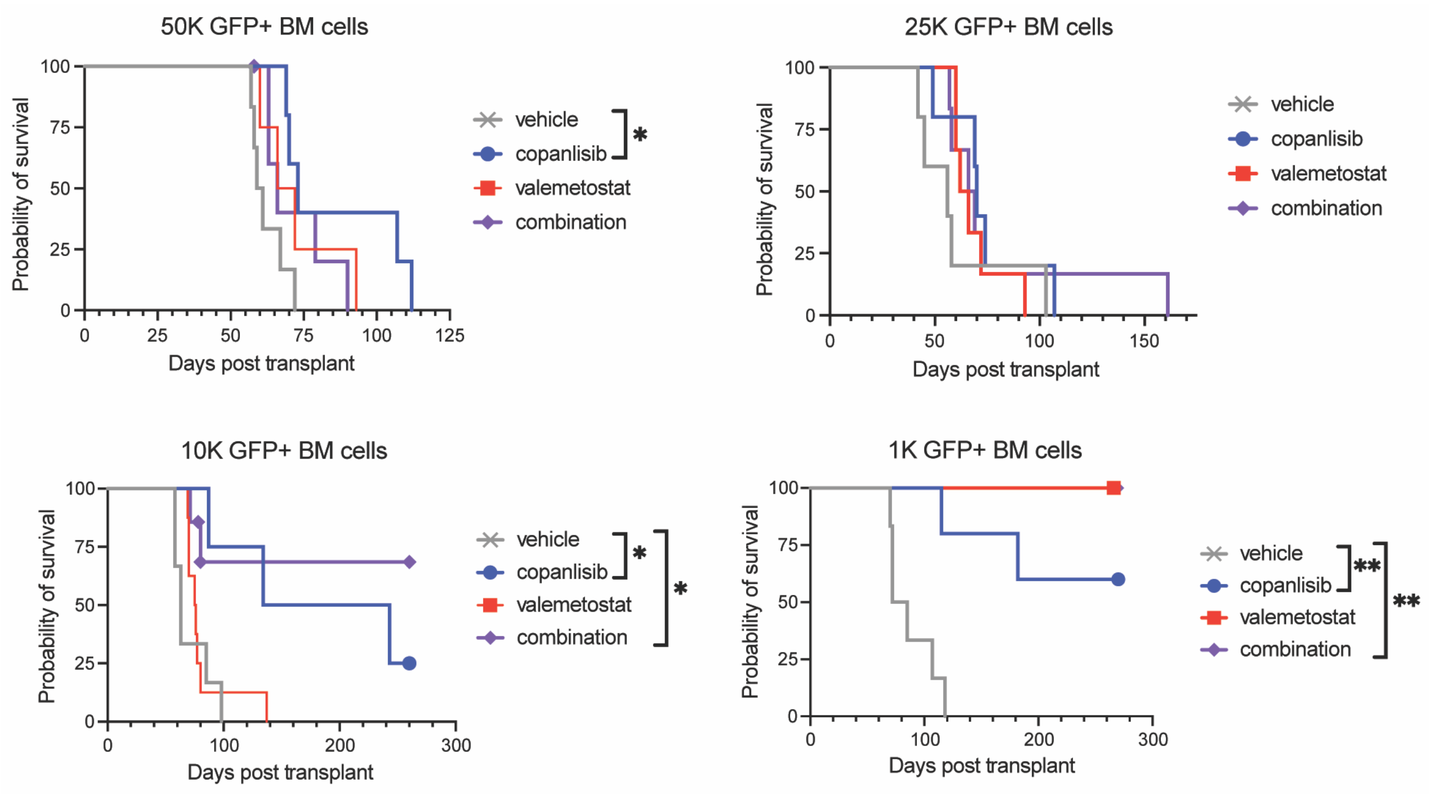


**Figure S12: PI3K Inhibition Cooperate with EZH1/2 Dual Inhibition to Target LSCs**
Kaplan-Meier survival curves of secondary transplant recipients injected with limiting numbers of KMT2A-MLLT3-GFP+ leukemic cells from primary transplant recipients after copanlisib +/- valemetostat treatment for 2 weeks in primary recipients only. Log-rank analysis was used. $**P \leq 0.01$, $*P \leq 0.05$

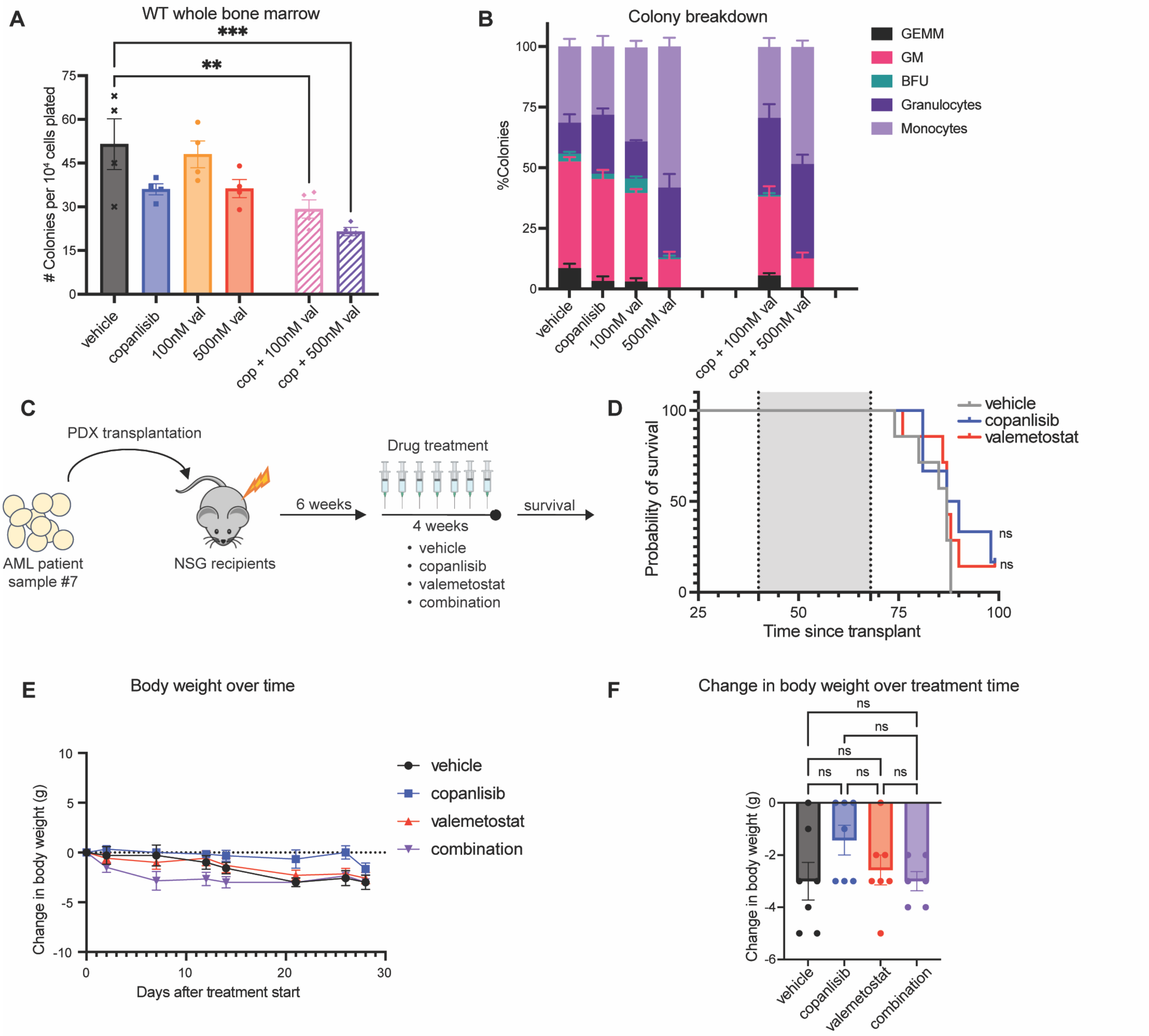


**Figure S13: There is a reasonable therapeutic window in healthy BM treated with copanlisib and valemetostat (A-B) (A)** Colony counts and **(B)** colony breakdown by morphology on colony assays performed on WT mouse BM cells with vehicle, copanlisib, valemetostat, or the combination **(C)** Schematic detailing PDX experimental plan. N=6 per group. **(D)** Line graph tracking the daily changes of body weight in grams of mice in each treatment group from PDX experiment. The gray shaded area represents the drug treatment duration. **(E)** Bar graph comparing net changes in body weight in grams of last day of treatment compared to first day. Each value is presented as mean +/- standard error of the mean (SEM). One-way ANOVA test with Tukey's multiple comparisons was used. ***P ≤ 0.001 **P ≤ 0.01 *P ≤ 0.05 ns ≥ 0.05.
